# Paradoxical mTORC1-Dependent microRNA-mediated Translation Repression in the Nucleus Accumbens of Mice Consuming Alcohol Attenuates Glycolysis

**DOI:** 10.1101/2023.11.29.569312

**Authors:** Yann Ehinger, Sophie Laguesse, Khanhky Phamluong, Alexandra Salvi, Yoshitaka J. Sei, Zachary W. Hoisington, Drishti Soneja, Sowmya Gunasekaran, Ken Nakamura, Dorit Ron

## Abstract

mTORC1 promotes protein translation, learning and memory, and neuroadaptations that underlie alcohol use and abuse. We report that activation of mTORC1 in the nucleus accumbens (NAc) of mice consuming alcohol promotes the translation of microRNA (miR) machinery components and the upregulation of microRNAs (miRs) expression including miR-34a-5p. In parallel, we detected a paradoxical mTORC1-dependent repression of translation of transcripts including Aldolase A, an essential glycolytic enzyme. We found that miR-34a-5p in the NAc targets Aldolase A for translation repression and promotes alcohol intake. Our data further suggest that glycolysis is inhibited in the NAc manifesting in an mTORC1-dependent attenuation of L-lactate, the end product of glycolysis. Finally, we show that systemic administration of L-lactate attenuates mouse excessive alcohol intake. Our data suggest that alcohol promotes paradoxical actions of mTORC1 on translation and glycolysis which in turn drive excessive alcohol use.

## Main

The mechanistic target of rapamycin (mTOR) is a ubiquitously expressed serine and threonine kinase that, when localized with specific accessory proteins, including the adaptor protein raptor, is termed mTORC1 (mTOR complex 1) ^1^. mTORC1 is activated by metabolism-relevant stimuli such as amino acids, lipid, oxygen, nutrients such as glucose as well as through the activation of growth factor signaling ^1,2^. mTORC1 plays a central role in cell growth by promoting the translation of a subset of mRNAs to proteins, as well as well as by enhancing lipid and nucleotide signaling ^1–3^. In contrast, mTORC1 also contributes to catabolism through the suppression of lysosomal biogenesis autophagy ^2^ and to aging ^1^. mTORC1 also plays a crucial role in sensing and regulating feeding and fasting processes ^1,3^. In the central nervous system, mTORC1 is known for its actions to promote the translation of dendritic spine proteins ^4,5^. Specifically, upon activation mTORC1 phosphorylates its substrates the p70 ribosomal S6 kinase (S6K) and the eukaryotic translation initiation factor 4E binding protein (4E-BP) ^6^. S6K then phosphorylates its substrate S6 ^6^, and these phosphorylation events promote the assembly of the translation initiation complex to initiate cap-dependent and independent mRNA translation of transcripts ^6^. mTORC1 together with its substrates are localized to ribosomes and the whole translation machinery is found in both cell body and dendrites ^6^. Not surprisingly, mTORC1 plays an important role in synaptic plasticity, learning and memory ^5,7,8^.

Because of its important role in cellular homeostasis, dysregulation of mTORC1 functions results in pathologies such as cancer, obesity, liver and pancreatic abnormalities ^1^ as well as neurodegenerative and neurodevelopmental diseases and psychiatric disorders ^5,9^ including addiction ^10^.

Research on the mechanism(s) by which mTORC1 drives adverse phenotypes associated with drugs of abuse has not been explored much. We found that the signaling cascade upstream of mTORC1 consisting of the small G protein H-Ras ^11^ and the kinases PI3K and AKT ^12^ is activated in the nucleus accumbens (NAc) of rodents consuming large quantities of alcohol. We further showed that pharmacological inhibition of H-Ras, PI3K, and AKT or knockdown of H-Ras in the NAc reduces excessive alcohol intake ^11,12^.

As mTORC1 is downstream of H-Ras/PI3K/AKT pathway ^13,14^, we tested whether chronic excessive alcohol intake activates mTORC1 in the NAc of rodents, and found that alcohol drinking produces a robust and long-lasting activation of mTORC1 signaling in the NAc^15^ and specifically in NAc shell ^16^. Using the selective mTORC1 inhibitor, rapamycin ^17^ we showed that mTORC1 in the NAc contributes to the development and/or maintenance of excessive alcohol use ^15,18–22^. To elucidate the mechanism(s) by which mTORC1 in the NAc drives excessive alcohol use, we conducted an RNAseq study and found that alcohol-mediated mTORC1 activation in the NAc of male mice led to the increase in the translation of 12 transcripts ^19^. Among the identified transcripts was the postsynaptic protein, Prosapip1 ^19^. We found that the translation of Prosapip1 is increased in response to alcohol-mediated mTORC1 activation, which in turn promotes the formation of F-actin, leading to synaptic and structural plasticity, alcohol self-administration and reward ^19^.

In addition, our RNAseq data suggested that alcohol-mediated activation of mTORC1 in the NAc also increases the translation of the microRNA machinery transcripts ^19^. The microRNA machinery is responsible for the repression of translation and mRNA degradation via the action of ∼22 nucleotides nucleotide sequences, microRNAs ^23,24^. After transcription by RNA polymerase II, pri-microRNAs are processed within the nucleus into pre-miRNAs by the microprocessor complex composed of Drosha and DGCR8 proteins ^25^. Once exported into the cytosol, the ribonuclease Dicer produces mature miRNAs that are then loaded into the miRNA-Induced Silencing Complex (RISC) ^25^. A perfect complementarity between a miRNA and its target mRNA leads to the cleavage of the target mRNA by miRISC resulting in mRNA degradation ^26^. An imperfect complementarity, however, results in reduced translation and/or stability of target mRNAs ^26^. Here, we examined the potential crosstalk between alcohol, mTORC1, and microRNA function.

## Results

### Alcohol increases the translation of Trax and GW182 in an mTORC1-dependent manner in the NAc

As mentioned above, using RNAseq analysis we previously identified the microRNA machinery components Trax, GW182 and CNOT4 as candidate transcripts whose translation was increased by alcohol in an mTORC1-dependent manner ^19^. To confirm the RNAseq data, male mice underwent intermittent access to 20% alcohol in a 2-bottle choice (IA20%2BC) paradigm for 7 weeks. This paradigm models humans who exhibit alcohol use disorder (AUD) ^27^. Three hours before the end of the last 24 hours alcohol withdrawal session, mice were systemically administered with vehicle or the selective mTORC1 inhibitor, rapamycin (20mg/kg). The NAc was removed 24 hours after the last alcohol withdrawal session, polysomes were purified to isolate actively translating mRNAs, and RT-qPCR analysis was conducted (**Fig. 1A, Supplementary Tables 1-2, alcohol drinking and statistical analysis)**. First, we replicated the published data ^19^ in a new cohort of animals and confirmed that the translating levels of Trax were indeed increased by alcohol and reduced back to baseline when mice were first pretreated with the mTORC1 inhibitor, rapamycin (**Fig. 1B**). Next we confirmed the RNAseq data using RT-qPCR and showed that the translation of GW182 is increased by alcohol in the NAc in an mTORC1-dependent manner (**Fig. 1C**). In contrast, the RNAseq data of CNOT4 in total NAc were not replicated (**Fig. 1D**). Since GAPDH is used as an internal control, we analyzed its level in response to alcohol and found no change in its translation in response to alcohol **(Supplementary Fig.1).**

**Figure 1.**
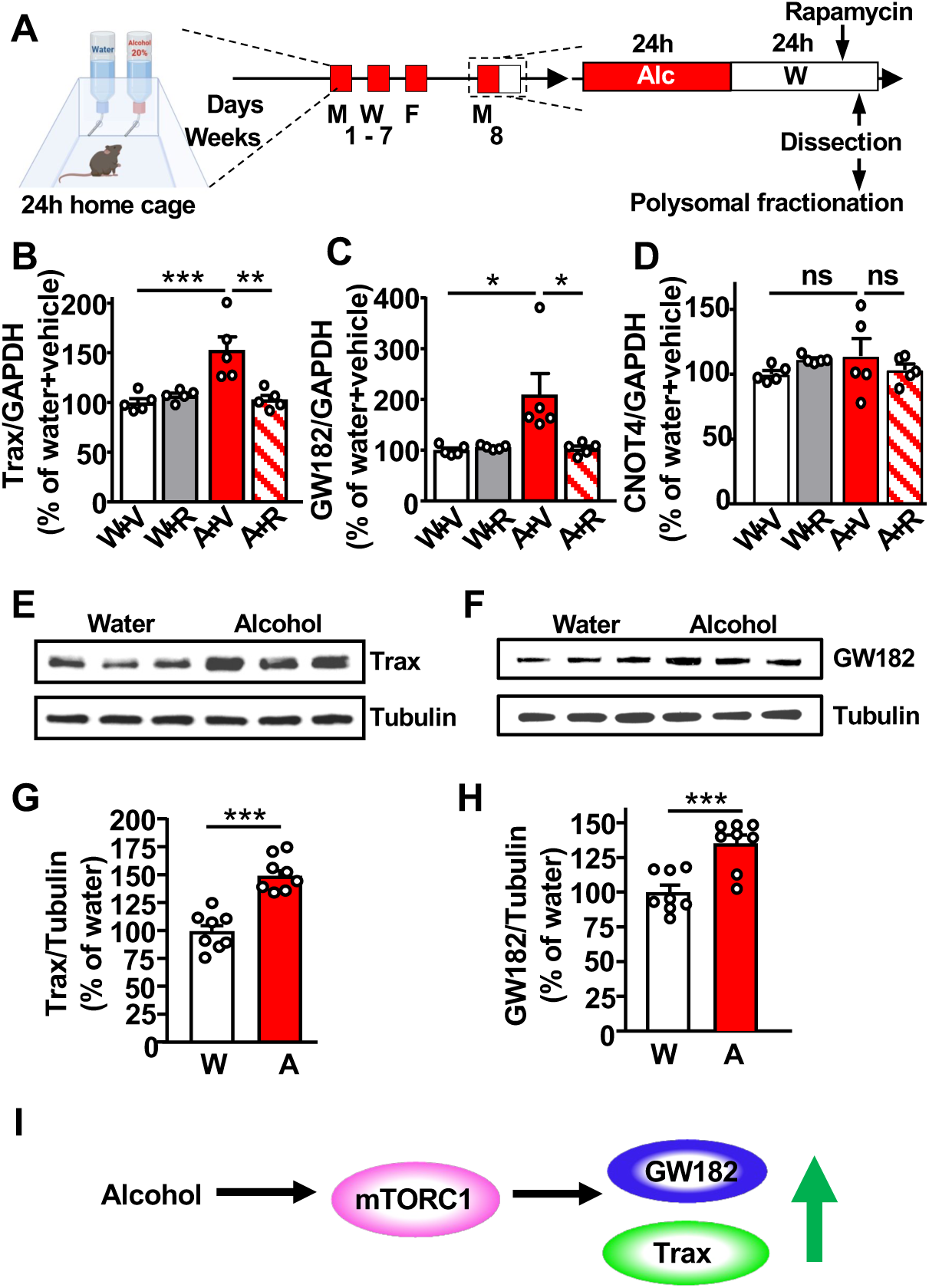
Alcohol via mTORC1 increases the translation of Trax and GW182 but not CNOT4 in the NAc. (**A**) Mice underwent 7 weeks of IA20%-2BC (**Supplementary Table 1**). Control animals had access to water bottles only. Three hours before the end of the last alcohol withdrawal period, mice were systemically injected with 20 mg/kg rapamycin or vehicle. The nucleus accumbens (NAc) was dissected from each of the four mouse groups (water+vehicle, water+rapamycin, alcohol+vehicle, alcohol+rapamycin) at the end of the last alcohol withdrawal period, were subjected to polysomal fractionation and RT-qPCR analysis. (**B-D**) Polysomal RNA levels of Trax (**B**), GW182 (**C**) and CNOT4 (**D**) were measured by RT-qPCR. Data are presented as the average ratio of each transcript to GAPDH±SEM and expressed as % of water+vehicle. Statistical analyses are depicted in **Supplementary Table 2**. *p<0.05, **p<0.01, ***p<0.001, ns: non-significant. **(E-H)** Trax (**E,G**) and GW182 (**F,H**) protein levels were determined by western blot analysis. ImageJ was used for optical density quantification. Data are presented as the average ratio of Trax or GW182 to Tubulin±SEM and are expressed as % of water control. Statistical analyses are depicted in **Supplementary Table 2**. ***p < 0.001. (**I**) Alcohol activates mTORC1 signaling in the NAc which in turn increases the translation of GW182 and Trax.

Next, a new cohort of mice that underwent 7 weeks of IA20%2BC was used to measure total mRNA of GW182 and Trax in alcohol and water drinking mice. We did not observe a change in the total amount of mRNA of either transcript (**Supplementary Fig. 2A-B, Supplementary Tables 1-2)**. Together, these data suggest that alcohol increases the translation but not the transcription and/or RNA stability of the microRNA machinery components GW182 and Trax.

Finally, we analyzed the protein levels of Trax and GW182 in water and alcohol drinking mice and found that the levels of both proteins were increased in the NAc of mice consuming alcohol as compared to water only drinking mice (**Fig. 1E-H, Supplementary Tables 1-2**). This increase was specific to the NAc as the levels of Trax and GW182 were not altered by alcohol in a neighboring striatal region, the dorsolateral striatum (DLS) (**Supplementary Fig. 2C-F, Supplementary Tables 1-2**). Together, these data suggest that chronic heavy alcohol use increases the levels of the microRNA machinery proteins Trax and GW182 in the NAc of drinking mice (**Fig. 1I**).

### Alcohol represses the translation of PPM1E, Aldolase A and Rbfox2 in an mTORC1-dependent manner in the NAc

The microRNA machinery is responsible for repression of translation or mRNA degradation ^23,24^. We speculated that alcohol-mediated increase in the translation of Trax and GW182 may point towards an unexpected link between mTORC1 and the microRNA machinery. Interestingly, while analyzing the RNAseq data ^19^, we observed that the translation of 32 transcripts was repressed by alcohol in an mTORC1-dependent manner (**Supplementary Table 3**). We classified the transcripts and found that 37% of them belong to the signal transduction category, 31% are part of the DNA/RNA machinery, 12% transcripts are associated with actin/cytoskeleton, and 12% with metabolic pathways **(Supplementary Fig. 3)**. We confirmed the RNAseq data of 3 random transcripts by measuring the mRNA levels in NAc polysomes of mice that underwent 7 weeks of IA20%2BC, and that were treated with vehicle or rapamycin (20 mg/kg) 3 hours before the last alcohol withdrawal session (**Fig. 2A-C, Supplementary Tables 1-2)**. We found that the translation of the glycolytic enzyme, Aldolase A^28^ (**Fig. 2A**), the serine/threonine Protein Phosphatase PPM1E (Mg2+/Mn2+ Dependent 1E) ^29^ (**Fig. 2B**), and the RNA binding protein Rbfox2 (RNA binding fox 1 homolog 2) ^30^ (**Fig. 2C**) was repressed by alcohol as compared to water only drinking mice. Importantly, we detected a reversal of translation repression in mice that were pre-treated with rapamycin (**Fig. 2A-C**). In contrast, we did not observe a change in the total mRNA levels of the transcripts suggesting that alcohol decreases the translation but not the transcription and/or mRNA stability of these 3 transcripts (**Supplementary Fig. 4A-C, Supplementary Tables 1-2**).

**Figure 2.**
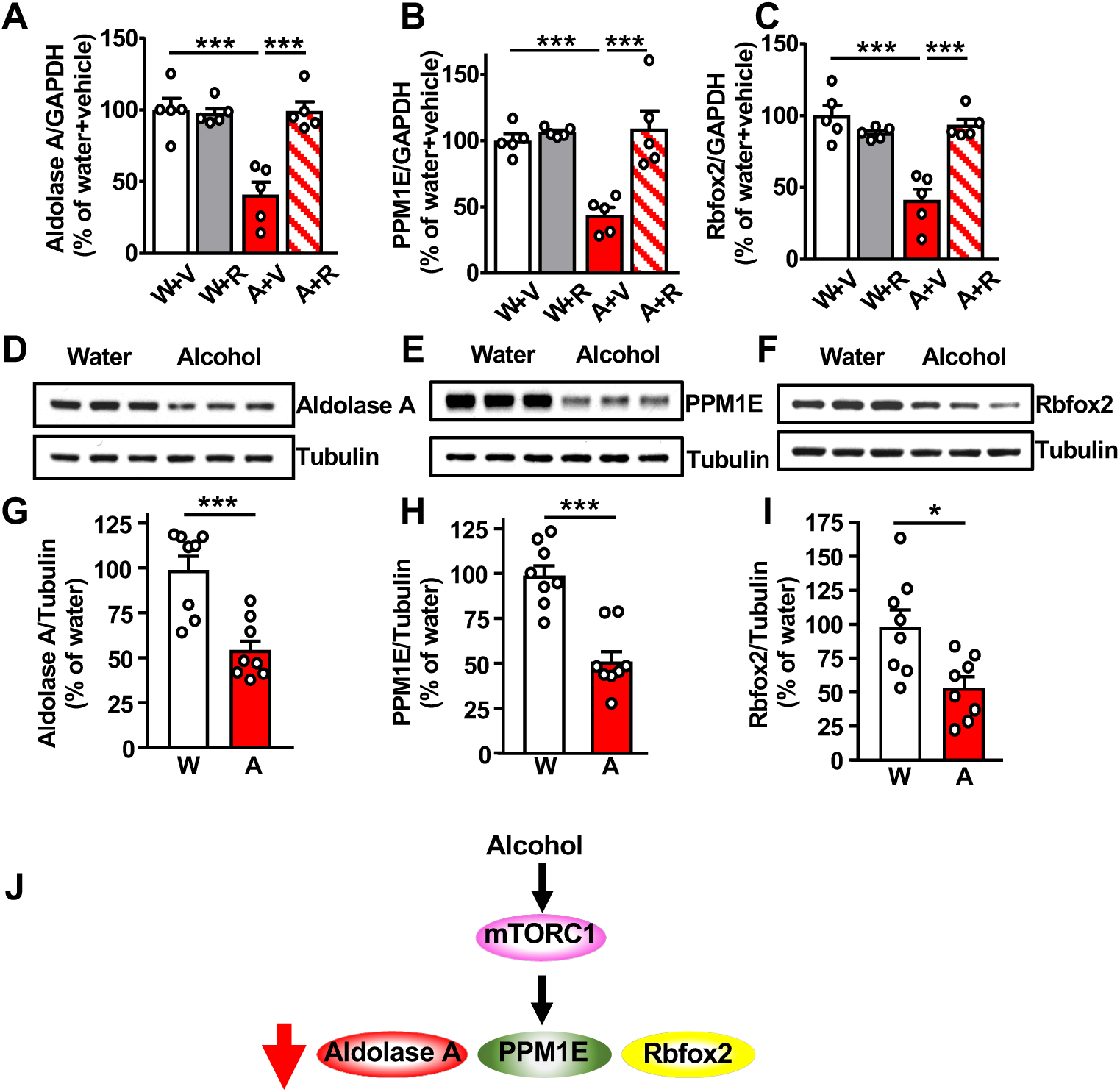
Alcohol via mTORC1 represses the translation of Aldolase A, PPM1E and Rbfox2 in the NAc. Mice underwent 7 weeks of IA20%-2BC (**Supplementary Table 1**) and were treated with 20mg/kg rapamycin as described above. (**A-C**) Polysomal mRNA levels of Aldolase A (**A**), PPM1E (**B**) and Rbfox2 (**C**). Data are presented as the average ratio of each transcript to GAPDH±SEM and expressed as the % of water+vehicle. Statistical analyses are depicted in **Supplementary Table 2**. ***p < 0.001. (**D-I**) Aldolase A (**D,G**), PPM1E (**E,H**) and Rbfox2 (**F,I**) protein levels were determined by western blot analysis. Data are presented as the average ratio of Aldolase A, PPM1E and Rbfox2 to Tubulin±SEM and are expressed as the % of water control. Statistical analyses are depicted in **Supplementary Table 2**. *p<0.05, ***p<0.001. (**J**) Alcohol activates mTORC1 signaling in the NAc which in turn decreases the translation of Aldolase A, Rbfox2, PPM1E.

Next, we analyzed the protein level of Aldolase A, PPM1E and Rbfox2 in water and alcohol drinking mice. We found that alcohol significantly decreases the level of the 3 proteins in the NAc (**Fig. 2D-I, Supplementary Tables 1-2**). The decrease was specific to the NAc, as the level of Aldolase A, PPM1E and Rbfox2 was not altered by alcohol in the DLS (**Supplementary Fig. 4D-I, Supplementary Tables 1-2**). Together, these data suggest that chronic heavy alcohol drinking increases the levels of the microRNA machinery proteins Trax and GW182 in the NAc and concomitantly represses the translation of Aldolase A, PPM1E and Rbfox2 (**Fig. 2J**).

### Alcohol-mediated mTORC1-dependent increase and decrease in translation is localized to D1 NAc neurons

The NAc is composed mainly of two neuronal subpopulations, dopamine D1 receptor (D1) and dopamine D2 receptor (D2) expressing medium spiny neurons (MSN) ^31^. We previously found that mTORC1 is specifically activated by alcohol in the shell but not the core of the NAc ^16^ and that the first drink of alcohol activates mTORC1 specifically in D1 neurons in the NAc shell ^20^. First, to determine if heavy chronic alcohol use activates mTORC1 in D1 NAc MSN, male D1-Cre mice were crossed with RiboTag mice in which GFP-fused ribosomal subunit RPL10 is expressed in the presence of Cre recombinase, enabling the detection of D1 neurons ^32^ (**Fig. 3A**). D1-Cre x RiboTag mice underwent 7 weeks of IA20%2BC and were sacrificed at the end of the last alcohol withdrawal session (**Fig. 3B, Supplementary Tables 1-2**). We found that alcohol significantly increases S6 phosphorylation in D1 neurons of the NAc shell (**Fig. 3C-D**) and that only 8.9% of phosphoS6 positive neurons are not D1 neurons (**Fig. 3E**). Together, these data suggest that the majority of mTORC1 activated by excessive alcohol use is localized to D1 NAc neurons.

**Figure 3.**
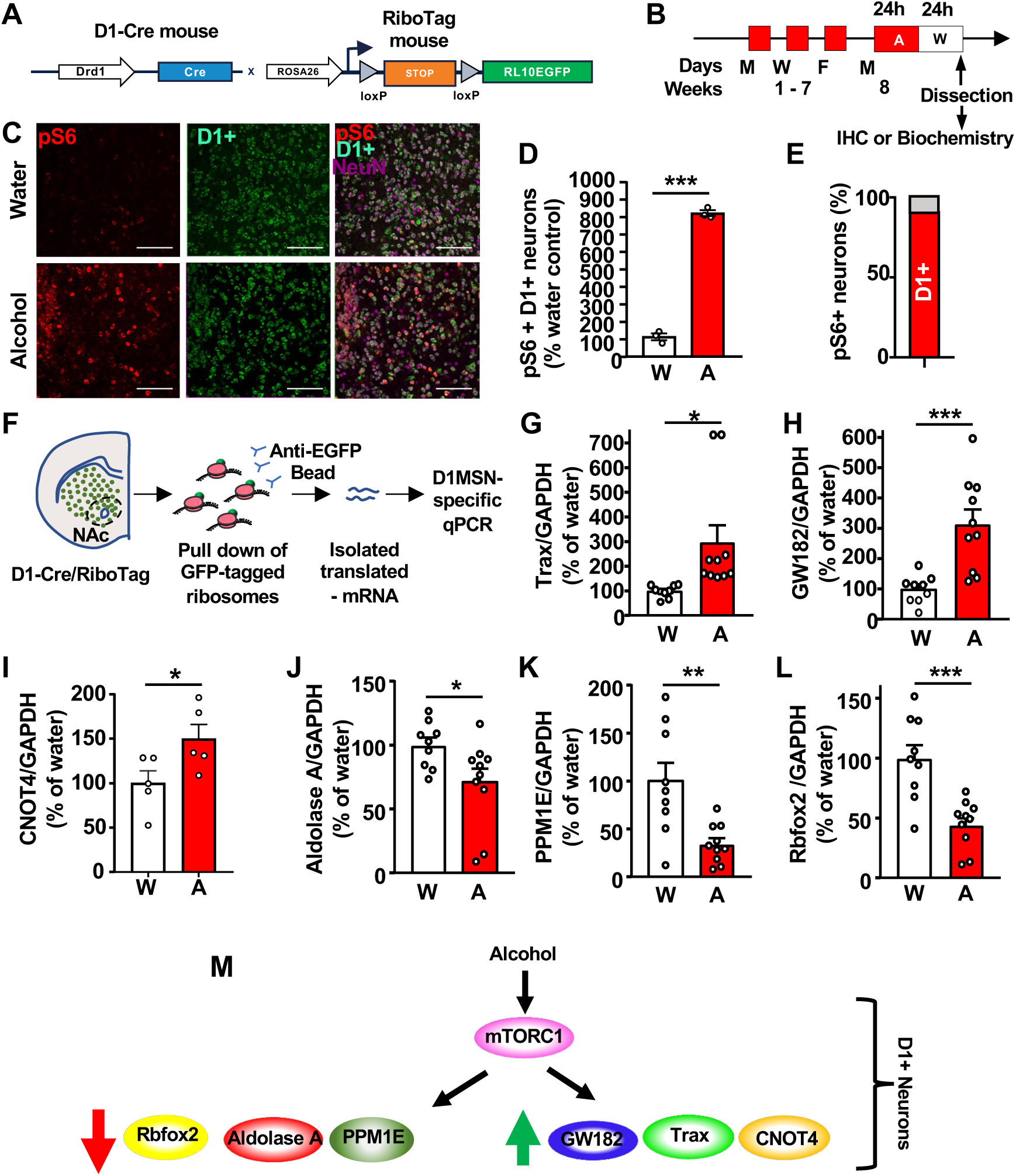
Alcohol activates mTORC1, increases the translation of Trax, GW182 and CNOT, and decreases the translation of Aldolase A, PPM1E, Rbfox2 in NAc D1+ neurons. (**A**) D1-Cre mice were crossed with RiboTag mice allowing the expression of RPL10-EGFP in D1-expressing neurons. (**B**) Mice underwent 7 weeks of IA20%2BC (**Supplementary Table 1**). Brains were dissected at the end of the last 24-hour withdrawal session as in Fig. 1A and processed for IHC or biochemistry analysis. (**C**) IHC analysis of phospho-S6 levels the NAc shell D1 neurons of drinking mice compared to water controls. Representative images of NAc at 20x magnification labeled with phospho-S6 in red, RPL10-GFP in green and NeuN in magenta. Bar scale 100µm. (**D**) Phospho-S6 and D1 labeled neurons are expressed as % of water controls. Statistical analyses are depicted in **Supplementary Table 2**. ***p<0.001. (**E**) Percentage of D1 positive vs. D1 negative NAc shell neurons labeled as phosphoS6 positive neurons. (**F**) Affinity purification of ribosomes from D1+ neurons by using anti-GFP magnetic beads followed by RNA isolation and RT-qPCR. (**G-L**) Polysomal mRNA levels in D1+ neurons of Trax (**G**), GW182 (**H**), CNOT4 (**I**), Aldolase A (**J**), PPM1E (**K**), Rbfox2 (**L**) after alcohol withdrawal were determined by RT-qPCR. Data are presented as the average ratio of a transcript to GAPDH±SEM and expressed as % of water control. Statistical analyses are depicted in **Supplementary Table 2**. *p<0.05, **p<0.01, ***p<0.001, ns: non-significant. (**M**) Alcohol activates mTORC1 signaling in D1+ NAc neurons which in turn increases the translation of GW182, Trax and CNOT4 and represses the translation of Aldolase A, Rbfox2 and PPM1E.

Next, we examined whether mTORC1-dependent alterations in translation are also localized to D1 NAc MSNs. To do so, we utilized D1-Cre x RiboTag mice which contain a GFP tag for affinity purification of D1 neuron ribosomes allowing for the isolation of mRNAs actively undergoing translation in D1 MSNs (**Fig. 3F**). D1-Cre x RiboTag mice underwent 7 weeks of IA20%2BC or water only, and mRNA levels were measured in the GFP immunoprecipitated fraction (**Fig. 3F, Supplementary Tables 1-2**). We detected an enrichment in Trax and GW182 translation in D1 NAc neurons (**Fig. 3G-H**). Interestingly, we also detected an increase in the level of CNOT4 mRNA undergoing translation in D1 MSNs (**Fig. 3I**). In parallel, translation of Aldolase A, PPM1E, and Rbfox2 was attenuated by alcohol in NAc D1 MSNs (**Fig. 3J-L)**. To test whether these increases and decreases in translation were specific to D1 NAc neurons, we utilized A2A-Cre (Cre expressed in D2 neurons) x RiboTag mice. mRNA levels were measured in the GFP immunoprecipitated fraction in water or alcohol drinking A2A-CrexRiboTag mice. We did not detect an increase in Trax, GW182 and CNOT translation in D2 NAc neurons **(Supplementary Fig. 5A-C).** In addition, PPM1E, Rbfox2, and Aldolase A levels were not decreased in D2 neurons **(Supplementary Fig. 5D-F).** Together, our data suggest that mTORC1-dependent translation activation and translation repression occur specifically in NAc D1 neurons (**Fig. 3M**).

### Identification of microRNAs targeting PPM1E, Aldolase A and Rbfox2

To determine whether miRs are responsible for the translation repression of PPM1E, Aldolase A and Rbfox2, we performed *in silico* analyses using algorithmic prediction tools miRwalk, miRDB and TargetScan to identify putative microRNAs targeting the 3 repressed transcripts. miR15b-5p, miR25-3p, miR-34a-5p and miR92a-3p were identified by all 3 prediction tools as potential specific miRs for PPM1E, Aldolase A, and Rbfox2 (**Fig. 4A**). We examine the level of the predicted miR in the NAc. Interestingly, we found that the level of all 4 miRs was elevated by alcohol in the NAc (**Fig. 4B, Supplementary Tables 1-2**). However, the expression of the miRs miR127-3p, miR122-5p, miR19b-3p, miR15a-5p, and miR-34a-3p were not altered by alcohol (**Fig. 4B, Supplementary Tables 1-2**). Since U6 was used as an internal control, we analyzed its level in response to alcohol and found no change **(Supplementary Fig. 6).** The alcohol-mediated increase in miR levels was specific to the NAc since the levels of these miRs were not altered by alcohol in the DLS (**Supplementary Fig. 7, Supplementary Tables 1-2**). Together, these data suggest the expression of miRs that are predicted to target Aldolase A, PPM2E and RbFox2 are increased in the NAc of mice that drink alcohol.

**Figure 4.**
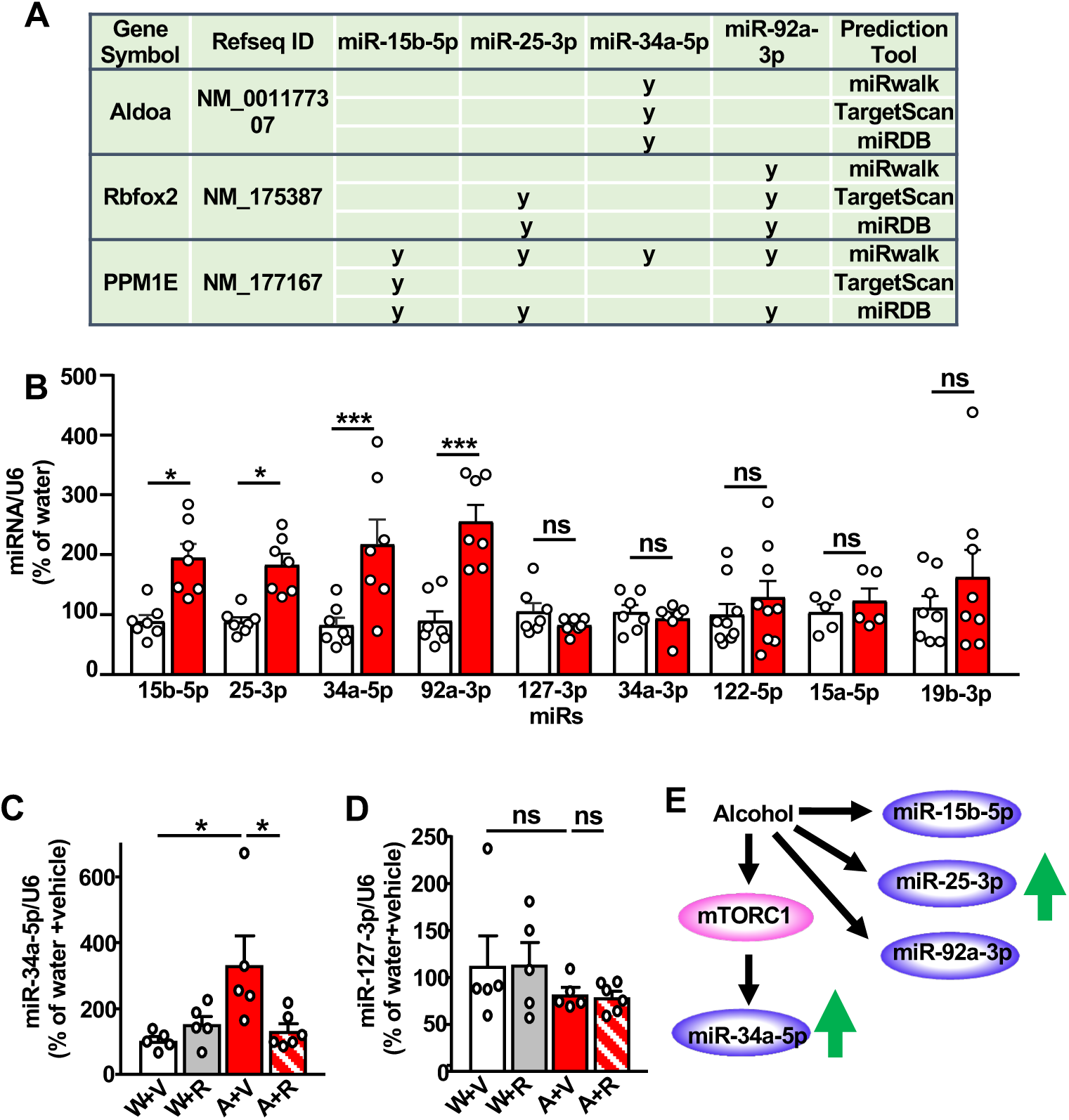
Identification of miR-15b-5p, miR-25-3p, miR-92-3p and miR-34a-5p which are increased by alcohol in the NAc; Activation of mTORC1 is required for alcohol-mediated increase of miR-34a-5p expression. **(A)** Potential miRNA-target interaction between miR-15b-5p, miR-25-3p, miR-92a-3p and miR-34a-5p the transcripts of interest: PPM1E, Aldolase A and Rbfox2. miRNA-target interactions were determined using miRWalk, TargetScan and miRDB. **(B)** Mice underwent 7 weeks of IA20%2BC or water only (**Supplementary Table 1**). Three hours before the end of the last 24 hours of alcohol withdrawal, the NAc was removed and the expression of miR-15b-5p, miR-25-3p, miR-34a-5p and miR-92-3p, as well as miR-127-3p and miR-34a-3p, miR-122-5p, miR-15a-5p, miR-19b-3p were measured by RT-qPCR. Statistical analyses are depicted in **Supplementary Table 2**. *p<0.05, ***p<0.001, ns: non-significant. **(C-D)** Mice underwent 7 weeks of IA20%2BC or water only (**Supplementary Table 1**). Three hours before the end of the last 24 hours of alcohol withdrawal session, mice that consumed alcohol (A) or water only (W) were injected with rapamycin (20mg/kg) (R) or vehicle (V). The levels of miR-34a-5p (**C**) and miR-127-3p (**D**) were measured by RT-qPCR. Statistical analyses are depicted in **Supplementary Table 2**. *p<0.05, ns: non-significant. (**E**) Alcohol increases the levels of miR-15b-5p, miR-25-3p, miR-92-3p and miR-34a-5p which are predicted to target the 3 transcripts shown in Fig. 3. Alcohol-mediated miR-34a-5p increase depends on mTORC1.

### miR-34a-5p targets Aldolase A in the NAc

We next focused on Aldolase A and its predicted miR, miR-34a-5p. First, we examined if the alcohol-mediated increase in miR-34a-5p expression in the NAc depends on mTORC1 and found that alcohol-mediated increase in miR-34a-5p expression is attenuated in mice that were pretreated with rapamycin (**Fig. 4C**, **Supplementary Tables 1-2**). In contrast, the level of miR127-3p was unaltered in mice drinking alcohol and treated with rapamycin (**Fig. 4D**, **Supplementary Tables 1-2**). We hypothesized that miR-34a-5p is required for the repression of Aldolase A translation and speculated that miR-34a-5p binds 3’UTR of Aldolase A. To test this possibility, we constructed luciferase expression plasmids containing Aldolase A 3’-UTR sequence or a mutant of the 3’UTR which does not contain the putative miR-34a-5p interaction site (**Fig. 5A-B**). Co-transfection of the miR-34a-5p with Aldolase A 3’-UTR plasmids into HEK293 cells significantly suppressed luciferase activity, while co-transfection of Aldolase A 3’-UTR with a control miR did not affect the luciferase activity (**Fig. 5C, Supplementary Table 2**). Mutation of the miR binding site within the Aldolase A 3’-UTR sequence abolished the effect of miR-34a-5p on the luciferase activity (**Fig. 5C, Supplementary Table 2**). These data demonstrate that Aldolase A 3’-UTR directly interacts with miR-34a-5p. To test whether miR-34a-5p is targeting Aldolase A *in vivo*, the NAc shell of mice was infected with a lentivirus expressing miR-34a (Ltv-miR-34a) (**Fig. 5D**). As shown in **Figure 5E-F (Supplementary Table 2)**, Aldolase A levels in the NAc were significantly reduced by miR-34a overexpression. Together, these data suggest that miR-34a is increased by alcohol via mTORC1 which in turn targets Aldolase A for translation repression.

**Figure 5.**
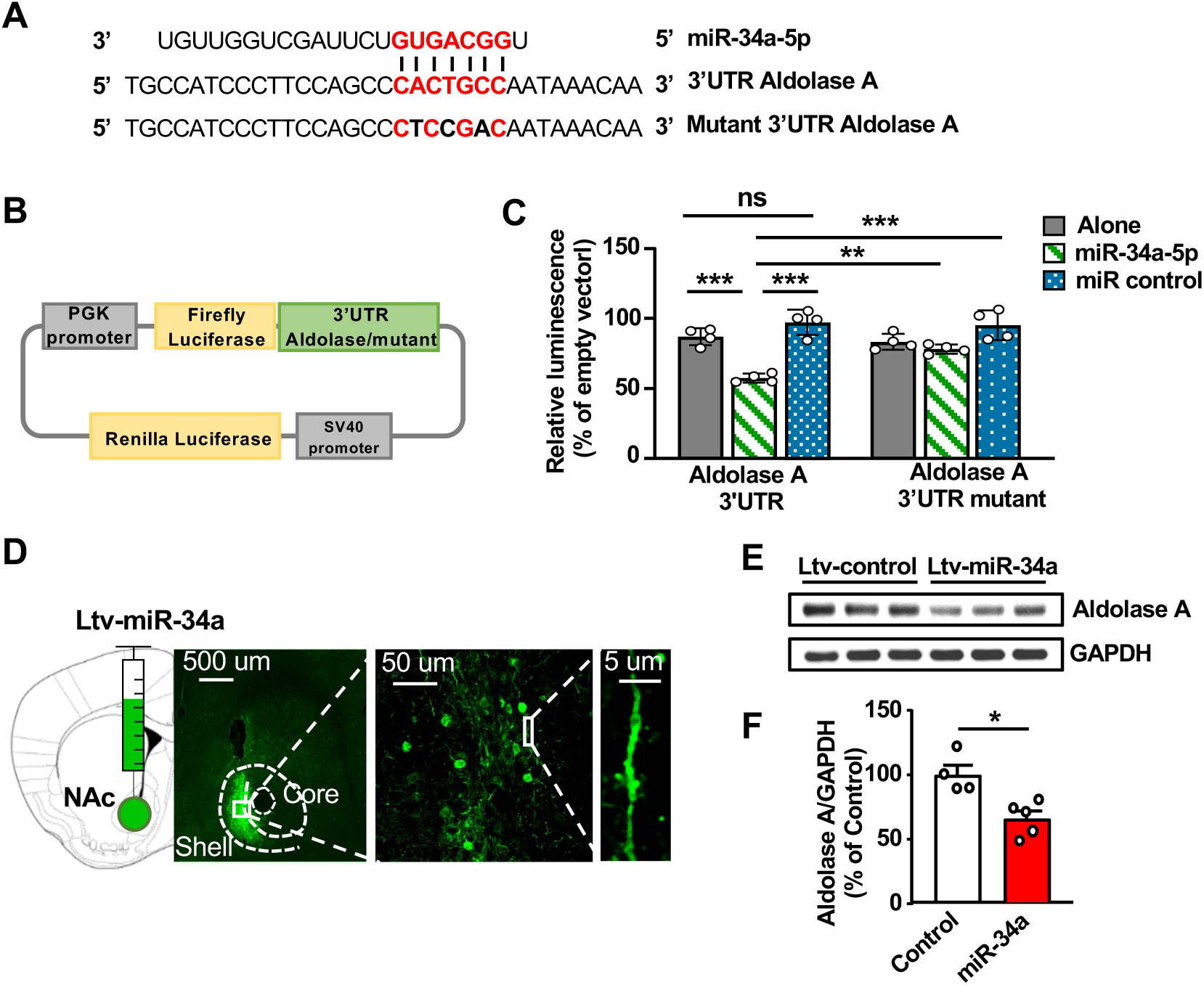
miR-34a-5p interacts with Aldolase A 3’UTR and represses Aldolase A levels in the NAc. (**A**) Predicted interaction sites (in red) of miR-34a-5p sequence (top) and Aldolase A 3’UTR (middle). Mutated Aldolase 3’UTR is shown at the bottom. (**B**) Map of Aldolase A 3’UTR or mutant 3’UTR cloned into a luciferase reporter vector. (**C**) HEK293 cells were co-transfected with a reporter vector containing Aldolase A 3’UTR or mutant 3’UTR and miR-34a-5p or a negative control. Bar graph depicts average ± SD expressed as Firefly luminescence normalized to Renilla luminescence and relative to a reference control. Statistical analyses are depicted in **Supplementary Table 2**. Gray bars: Aldolase A or mutant Aldolase A. Hatched green bars: Aldolase A or mutant Aldolase A + miR-34a5p. Dotted blue bars: Aldolase A or mutant Aldolase A + miR control. **p<0.01, ***p<0.001, ns: non-significant. (**D**) Lenti-miR-34a-GFP infected NAc neurons. (**E-F**) miR-34a-5p or GFP was expressed in the NAc and Aldolase A protein level was evaluated by western blot analysis, quantified as a ratio of Aldolase A/GAPDH and depicted as % of Aldolase A levels in the NAc of GFP infected mice. Statistical analyses are depicted in **Supplementary Table 2**. *p<0.05.

### Alcohol reduces glycolysis in the NAc via mTORC1

Aldolase A belongs to the family of the Fructose Diphosphate Aldolase enzymes ^28^. Aldolase A is highly expressed in neurons and is the major isoform expressed in the striatum ^33,34^. Aldolase C is also expressed in the brain ^35^, however we found that Aldolase C protein levels in the NAc were unaltered by alcohol (**Supplementary Fig. 8**). Aldolase A is a critical enzyme in glycolysis, a ten-step metabolic pathway resulting in the production of lactate ^36^, and ATP molecules through the tricarboxylic acid (TCA) cycle in the mitochondria ^37^. Aldolase A catalyzes the conversion of Fructose 1,6-Bisphosphate (F1,6BP) to Glyceraldehyde 3-phosphate (G3P) and dihydroxyacetone phosphate (DHAP) ^37^ (**Fig. 6A**). Since alcohol represses Aldolase A translation and protein levels in the NAc, we hypothesized that alcohol-mediated activation of mTORC1 leads to a reduction in glycolysis. To examine the possibility, we first conducted a metabolomics study in which we used mice that underwent IA20%2BC for 7 weeks (**Supplementary Tables 1**). Twenty-four hours after the last drinking session, the NAc was dissected followed by metabolites extraction and mass spectrometry analysis. We found that the end product of glycolysis, lactate ^36^, as well as other metabolites within the TCA cycle such as citrate, a-ketoglutarate and malate, were reduced by alcohol (**Fig. 6B, Supplementary Table 2**), suggesting that alcohol reduces glycolysis in the NAc. The attenuation of glycolysis was also not due to changes in the expression of the main neuronal and astrocyte glucose or lactate transporters (**Supplementary Fig. 9**) ^19^. Ter Horst reported that D1 MSNs in the NAc of mice regulate glucose tolerance sensitivity in the periphery ^38^. We therefore investigated whether glucose tolerance was affected in alcohol drinking mice. As shown in **Supplementary Figure 10**, blood glucose level during the glucose tolerance test was not altered by 4 or 7 weeks of IA20%2BC, suggesting that alcohol’s effect on glucose metabolism is centrally localized.

**Figure 6.**
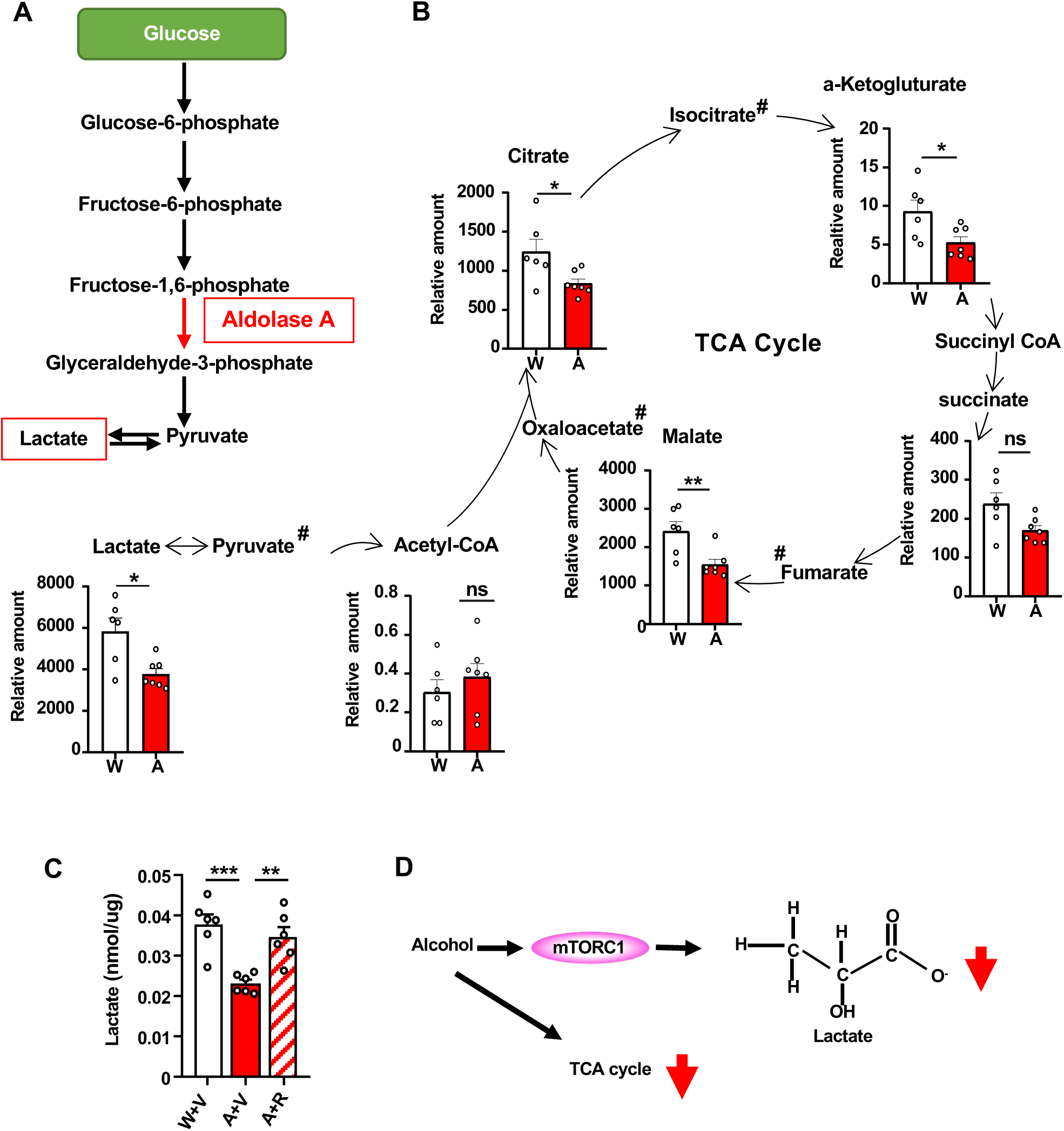
Alcohol decreases TCA cycle metabolites, including lactate which depends on mTORC1. (**A**) Aldolase A in the glycolysis pathway converts F1,6BP to G3P and DHAP. Lactate is the final product of glycolysis. (**B**) After 7 weeks of IA20%2BC (**Supplementary Table 1**) or water only, the NAc was dissected after 24 hours of alcohol withdrawal and metabolite levels were measured. Data are presented as relative amount of individual metabolites in water vs. alcohol. Statistical analyses are depicted in **Supplementary Table 2**. *p<0.05, **p<0.01, ns: non-significant. # metabolites that were not included in the panel. (**C**) Mice underwent 7 weeks of IA20%2BC (**Supplementary Table 1**) or water only. Three hours before the end of the last alcohol withdrawal period, mice were systemically injected with 20mg/kg rapamycin or vehicle. The NAc was dissected after 3 hours, and lactate level was measured using a colorimetric assay. Statistical analyses are depicted in **Supplementary Table 2**. **p<0.01, ***p<0.001. (**D**) Alcohol reduces TCA metabolites in the NAc and lactate which depends on mTORC1.

To examine if alcohol-mediated reduction in lactate content identified in the metabolomics study is mTORC1-dependent, mice underwent 7 weeks of IA20%2BC (**Supplementary Table 1)**. Three hours before the end of the last withdrawal session, mice were treated with vehicle or rapamycin. We found that lactate content in the NAc was significantly decreased by alcohol in mice pretreated with vehicle (**Fig. 6C, Supplementary Table 2**). In contrast, lactate content in the NAc was back to basal levels in mice that consumed alcohol and that were pretreated with rapamycin (**Fig. 6C, Supplementary Table 2**). Together, these data suggest that a consequence of the mTORC1↑/miR-34a-5p↑/Aldolase A↓ pathway is the attenuation of glycolysis and the reduction in lactate content in the NAc (**Fig. 6D**).

### Overexpression of miR-34a-5p in NAc D1 neurons accelerates heavy alcohol intake

We previously found that mTORC1 in the NAc plays an important role in mechanisms underlying alcohol drinking behaviors ^15,18,19^. Since we found that the activation of mTORC1 in D1 NAc neurons by alcohol promotes the increase in miR34a-5p expression leading to the attenuation of Aldolase A translation in D1 neurons, we speculated that the mTORC1↑/miR-34a-5p↑/Aldolase A↓ pathway in the NAc D1 MSNs contributes to the development of excessive alcohol consumption. We further reasoned that if this pathway contributes to neuroadaptations that drive alcohol intake, then overexpression of miR-34a-5p in NAc D1 MSNs will repress the translation of Aldolase A, and increase alcohol consumption. To test this, the NAc shell of D1-Cre mice was infected with AAV2-Flex-miR-34a or an AAV2-Flex-control virus that only contains a fluorescent reporter (**Supplementary Fig. 11**). Three weeks later, mice were subjected to IA20%2BC (**Timeline Figure 7A**). We found that overexpression of miR-34a-5p, specifically in D1 MSNs, increased and accelerated alcohol drinking as compared to mice infected with AAV2-Flex-control (**Fig. 7B, Supplementary Table 2**) which was not due to increased locomotive activity (**Fig. 7C**).

**Figure 7.**
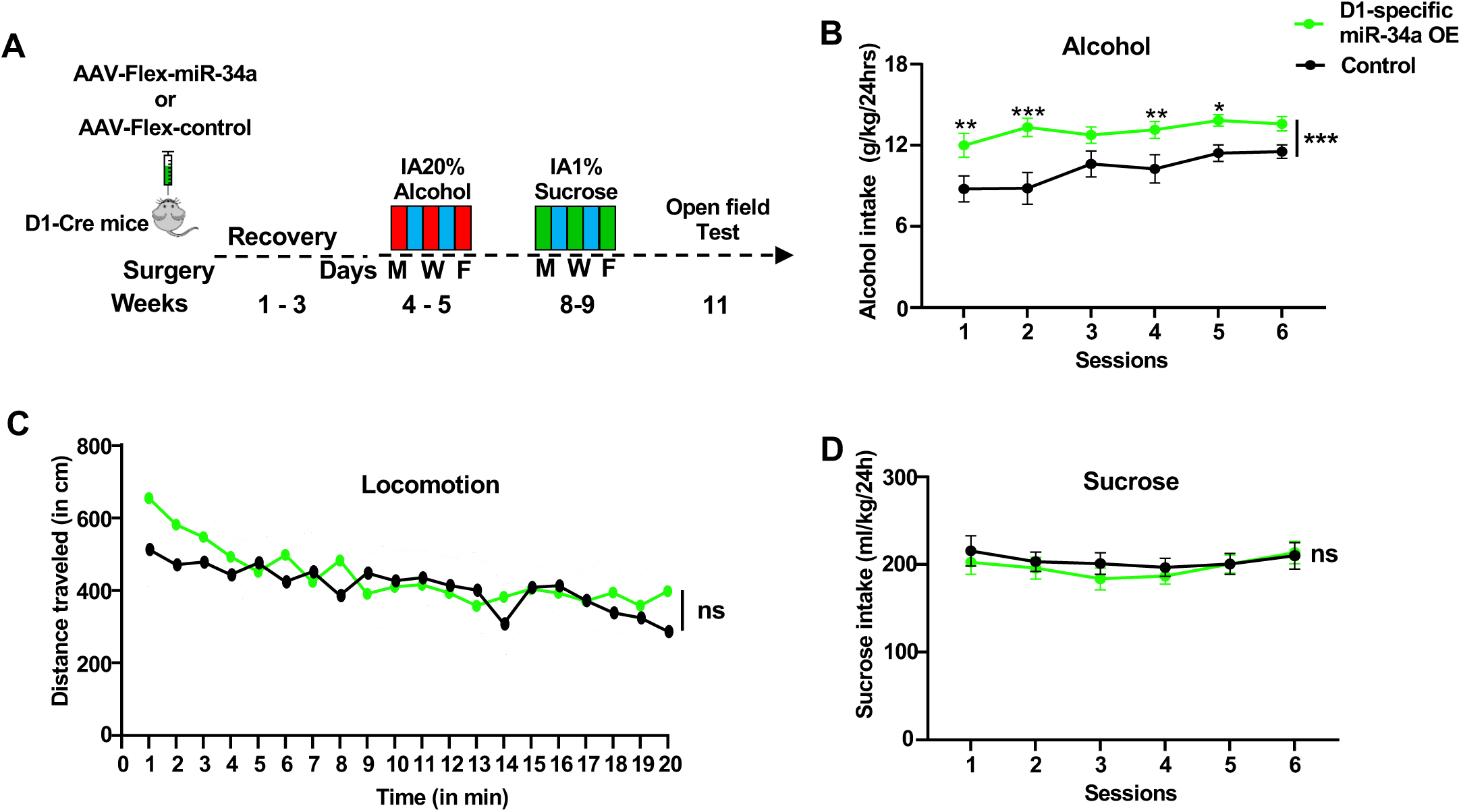
Overexpression of miR-34a in NAc D1 neurons increases alcohol consumption but does not affect locomotion or sucrose intake. **(A)** Experimental timeline. Mice received a bilateral infusion of AAV2-Flex-miR-34a or AAV2-Flex-control in the NAc shell of D1-Cre mice. Three weeks after surgery, mice underwent 2 weeks of IA20%2BC followed by 2 weeks of water only and 2 weeks of IA1%Sucrose2BC. Mice were then subjected to the open field test. **(B)** The NAc of D1-Cre mice was infected with AAV2-Flex-control or AAV2-Flex-miR-34a-GFP. Three weeks later, mice underwent IA20%2BC, and alcohol consumption was measured daily for 6 sessions. Statistical analyses are depicted in Supplementary Table 2. *p<0.05, **p<0.01, ***p<0.001. **(C)** The NAc of D1-Cre mice was infected with AAV2-Flex-control or AAV2-Flex-miR-34a. Mice were placed in an open field and mice movement was recorded for 20 minutes. Locomotion is depicted in 1-minute bins. Statistical analyses are depicted in Supplementary Table 2. ns: non-significant. **(D)** AAV2-Flex-control or AAV2-Flex-miR-34a-GFP infected mice underwent 2 weeks of IA1%Sucrose2BC. Statistical analyses are depicted in Supplementary Table 2. ns: non-significant.

Alcohol often acts distinctly from natural reward ^39,40^, and we previously showed that the mTORC1 signaling in the NAc is not activated in response to sucrose intake ^16^. We further discovered that inhibition of mTORC1 by rapamycin does not alter sucrose intake ^15^. We hypothesized that mTORC1↑/miR-34a-5p↑/Aldolase A↓ pathway in D1 MSNs does not play a role in sucrose consumption. To test this hypothesis, D1-Cre mice infected with AAV2-Flex-miR-34a or AAV2-Flex-control were subjected to IA 1% sucrose 2BC (**Timeline Figure 7A**). As shown in **Figure 7D**, the effect of D1-specific overexpression of miR-34a was specific for alcohol as it did not affect sucrose consumption.

### L-Lactate reduces heavy alcohol intake

As depicted above the activation of mTORC1↑/miR34a-5p↑/ Aldolase A↓ pathway results in a decrease in glycolysis and its final product lactate. We speculated that replenishing lactate levels will attenuate alcohol consumption. To test this hypothesis, mice underwent 7 weeks of IA20%2BC (**Supplementary Table 1**) and were then treated with L-lactate (2g/kg) supplement, which was administered subcutaneously (s.c.) 30 minutes before a drinking session, and alcohol and water intake were measured. We found that systemic administration of L-lactate was sufficient to reduce binge alcohol intake (**Fig. 8A, Supplementary Table 2**). At the 4 hours time points, in parralel to the reduction of alcohol intake, a concominent increase in water (**Fig.8B**) and total fluid (**Fig. 8C**) were detected. However, alcohol and water intake at the end of the 24-hour drinking session were not affected by L-lactate (**Fig. 8D-F**). Recently Lund et al. reported that the high amount of sodium in the L-lactate solution induces dehydration and significantly increases water intake ^41^ as detected in **Figure 7B-C, Supplementary Table 2**. To rule out the possible adverse effect of sodium on alcohol drinking, a solution of sodium chloride (NaCl iso-osmolar solution, 1g/kg) was administered 30 minutes before a drinking session, and alcohol and water intake were measured. NaCl-treated mice showed a significant increase in water consumption, whereas their alcohol intake was unchanged (**Supplementary Fig. 12, Supplementary Table 2**). These results suggest that the decrease in alcohol consumption by L-lactate administration was driven by L-lactate and not by sodium.

**Figure 8.**
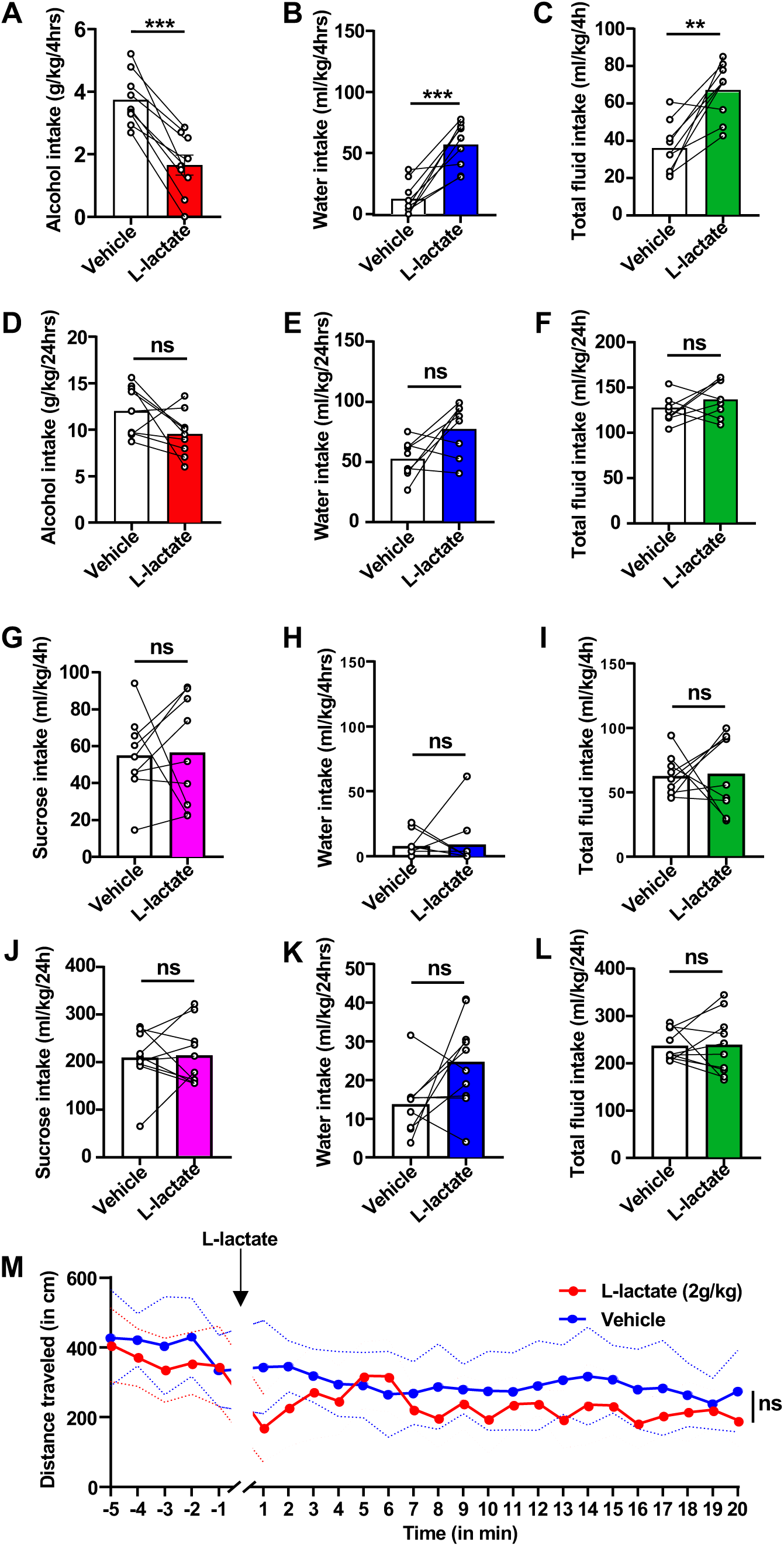
Subcunenious administration of L-lactate attenuates alcohol consumption but does not affect locomotion or sucrose intake. (**A-F**) Mice underwent 7 weeks of IA20%2BC (**Supplementary Table 1**). Mice received a single administration of L-lactate (s.c. 2g/kg) or PBS on weeks 8^th^ and 9^th^ 30 minutes before the beginning of the 24 hours alcohol drinking session in a counterbalanced manner. Alcohol and water were measured at the 4 hours (**A-C**) and 24 hours (**D-F**) time points. Statistical analyses are depicted in **Supplementary Table 2**. **p<0.01, ***p<0.001, ns: non-significant. (**G-L**) Mice underwent 2 weeks of IA1%Sucrose2BC. Control animals had access to water only. On weeks 3 and 4, mice were s.c. injected with L-lactate (2g/kg) or PBS in a counterbalanced manner 30 minutes before the beginning of a 24-hour drinking session. Sucrose and water intake were measured 4 hours (**G-I**) and 24 hours (**J-L**) later. Statistical analyses are depicted in **Supplementary Table 2**. ns: nonsignificant. **(M)** Mice were habituated for 5 minutes in the open field apparatus. Mice were then injected s.c. with 2g/kg L-lactate or PBS before being placed back in the open field and movement was recorded for an additional 20 minutes. Locomotion is depicted in 1-minute bins. Statistical analyses are depicted in **Supplementary Table 2**. ns: non-significant.

We next assessed whether the L-lactate effect is specific to alcohol or is shared with other rewarding substances. A new cohort of mice underwent 2 weeks of IA 1% sucrose 2BC and was then treated with L-lactate (2g/kg) subcutaneously 30 minutes before a drinking session. We found that L-lactate administration did not alter sucrose intake (**Fig. 8G-L**). Finally, we did not detect alteration in mice locomotion following the administration of L-lactate (**Fig. 8M**). Together, our data imply that the signaling cascade that includes miR-34a-5p and lactate drives alcohol intake (**Fig. 9**).

**Figure 9.**
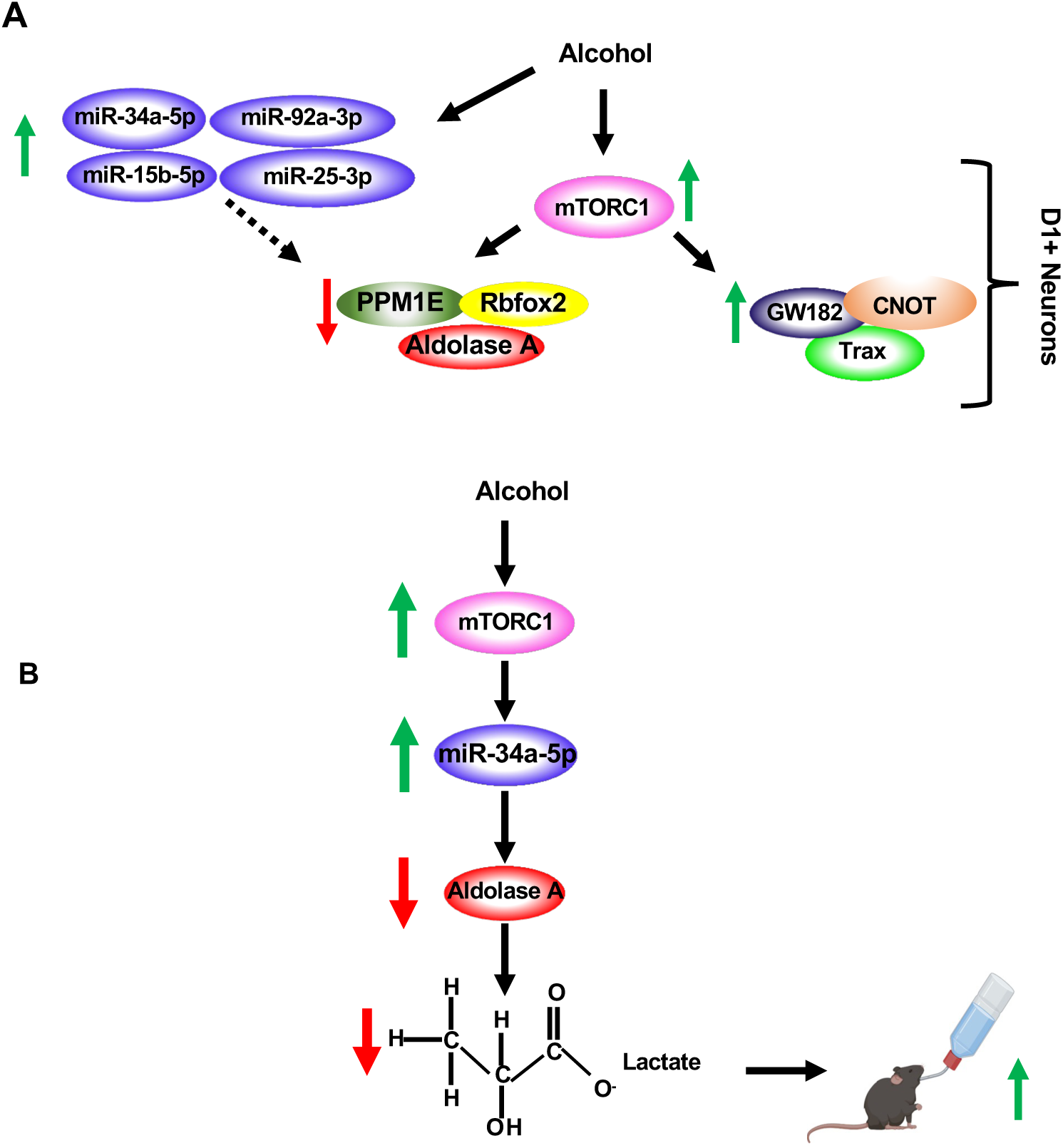
(**A**) Alcohol activates mTORC1 signaling in D1+ NAc neurons which in turn increases the translation of GW182, Trax and CNOT4 and represses the translation of Aldolase A, Rbfox2 and PPM1E. In parallel, alcohol increases the levels of miR-15b-5p, miR-25-3p, miR-92-3p and miR-34a-5p which are predicted to target Aldolase A, Rbfox2 and PPM1E. (**B**) Alcohol activates mTORC1 signaling in the NAc which increases the level of miR-34a-5p, repressing the translation of Aldolase A and decreasing the level of L-lactate, promoting further drinking.

## Discussion

We found that alcohol, by activating mTORC1 in the NAc shell D1 neurons, increases the translation of microRNA machinery transcripts and the levels of miRs, including miR-34a-5p. Alcohol-dependent activation of mTORC1 also represses the translation of a number of transcripts including Aldolase A specifically in D1 NAc neurons. We further showed that miR-34a-5p targets Aldolase A for translation repression. As a result of alcohol-mediated mTORC1-dependent reduction in Aldolase A levels in the NAc, glycolysis is inhibited, and lactate levels are reduced. Our results therefore suggest that the pathway mTORC1↑/miR-34a-5p↑/Aldolase A↓/glycolysis↓Lactate↓ in the NAc contributes to escalation of alcohol intake (**Fig. 9**).

Our study focused on male mice as mTORC1 is not activated after binge drinking or withdrawal in the NAc of female mice ^42,43^. Along with these findings, we found that systemic administration of rapamycin in female mice, unlike in male mice, does not reduce alcohol consumption ^43^. Similarily, Cozzoli et al. reported that intra-NAc administration of rapamycin does not change alcohol intake in female mice ^42^. What could be the root cause for the sex-specific effects of alcohol on mTORC1 signaling? One possibility is that the localization of components in the mTORC1 signaling cascade in the NAc is *a priori* different in male vs. female mice. It is also plausible that stress ^44^, and/or sex hormones ^45^, and/or sex-chromosome components ^46,47^ interact with alcohol and influences the activation of mTORC1 signaling differently in the NAc of male and female mice. Together, our data suggest a sex-specific interaction between alcohol and mTORC1, which is currently being investigated.

We previously showed that both binge alcohol drinking and withdrawal activate mTORC1 in the NAc ^15,16^. Here we show that mTORC1 activation, as well as translation enhancement and repression by alcohol, are localized to D1 neurons. What could be the mechanism for alcohol-mediated mTORC1 activation in NAc D1 neurons? Withdrawal from alcohol drinking increases the glutamatergic tone in the brain ^48^ and specifically in the NAc ^49^. Stimulation of glutamate metabotropic mGlu1/5 receptors activates mTORC1 in hippocampal neurons ^50^, and we previously found that stimulation of the NMDA receptor (NMDARs) activates mTORC1 in the orbitofrontal cortex (OFC) ^51^. Thus, it is plausible that the release of glutamate by cortical inputs specifically onto D1-MSN triggers mTORC1 activation in NAc D1 MSNs and the initiation of the cascade. We also found that mTORC1 is activated in the NAc during a short binge drinking session ^15,16^, a time point in which the NMDARs are inhibited ^52^. Actutely, alcohol was shown to block the NMDARs in the hippocampus ^53^, and we previously found that acute alcohol exposure produces a robust inhibition of the activity of NMDARs in the NAc ^52^. Curiously, inhibition of the NMDARs was shown to activate mTORC1 in the prefrontal cortex (PFC) ^54^ and in primary hippocampal neurons ^55^. Furthermore, the NMDAR inhibitor and antidepressant drug, ketamine, activates mTORC1 ^56^. Therefore, it is plausible that during binge drinking of alcohol, mTORC1 is activated by NMDAR inhibition and that the activation is maintained via glutamate binding to the NMDAR and/or mGlu1/5 during withdrawal.

RNAseq analysis aimed to identify alcohol-mediated mTORC1-dependent translatome suggested that alcohol enhances the translation of miR machinery proteins CNOT4, Trax and GW182 ^19^. We found that Trax and GW182 translation was increased in the NAc and specifically in NAc shell D1 neurons of mice consuming alcohol. Interestingly, alcohol-mediated translation of CNOT4 was only detected in D1 neurons which could be due to CNOT4 signal being lost upon harvesting the whole NAc. Together, these data suggest that mTORC1 activation by alcohol upregulates the translation of Trax, GW182 and CNOT4 in D1 NAc shell neurons. GW182 is an essential part of the translation repression machinery ^24,25^. Trax and its binding partner Translin were reported to promote mRNA degradation ^57,58^. Chen et al. found that the mRNA silencing CCR4-NOT complex, which CNOT4 is part of, hooks onto GW182 and recruits DDX6 to repress miR-target mRNAs ^59^. In parallel to the increase in the translation of the microRNA machinery transcripts, we identified alcohol-mediated mTORC1-dependent translation repression as well as the elevation of levels of specific miRs. Interestingly, hyperactivation of mTORC1 in cells results in an increase in microprocessor activity, a nuclear complex including Drosha and its partner DGCR8, thereby promoting miR biogenesis ^60^. Interestingly, excessive mTORC1 activation in tuberous sclerosis complex (TSC) disorder and Alzheimer’s disease leads to the reduction of specific proteins ^61–63^. Further studies are required to test the direct link between the upregulation of the microRNA machinery proteins GW182, Trax, and CNOT4 and repression of translation by alcohol and whether the increase in these transcripts promotes, as we predict, the function and/or efficacy of the microRNA machinery.

Our RNAseq data suggest that one of the consequences of mTORC1 activation by alcohol in the NAc is the repression of translation of 32 transcripts. These data are unexpected as mTORC1’s main role is to promote translation ^4,8^, whereas miRs function is to degrade mRNAs and to repress translation ^23^. Previous data suggest that miRs target components of mTORC1 signaling and thus inhibit mTORC1 function ^64–66^. For instance, miR199a-3p and miR100 directly target mTOR itself ^64,65^, and activation of mTORC1 by high level of nutrients represses miRs biosynthesis by the mRNA degradation of Drosha ^66^. Thus, the fact that mTORC1 activation under certain circumstances, e.g., heavy alcohol use promotes miRNA biosynthesis and represses the translation of numerous transcripts challenges the conventional notion and suggests context-specific effects of mTORC1 signaling on miR expression.

Interestingly, we observed that in parallel to the repression of translation, alcohol increases the levels of miRs that were identified by bioinformatic tools to potentially target the 3 tested transcripts. Importantly, we show that alcohol-mediated biosynthesis of at least one of the miRs (miR-34a-5p) depends on mTORC1, thus directly linking mTORC1 to the microRNA machinery. The miR34 family has been associated with neurodegenerative disease ^67^, psychiatric disorders ^67–73^ as well as fear, anxiety, stress ^67,70,73^, memory deficits and cognition ^71,72^. Here, we identified new roles for miR-34a-5p that are localized to the NAc, e.g., repression of Aldolase A translation and upregulation of alcohol drinking. Interestingly, mTORC1 in TSC2-deficient cells has been shown to increase the level of miR21, in turn promoting mTORC1-driven tumorigenesis ^63^. Raab-Graham and colleagues previously reported that inhibition of rapamycin increased Kv1.1 synaptic levels in hippocampal dendrites ^74^ through miR129 ^75,76^, although in contrast to our study, miR129 expression was mTORC1-independent ^75,76^.

We found that alcohol represses the translation of Aldolase A in the NAc. Aldolase A is an essential glycolytic enzyme that catalyzes the conversion of F1,6BP to G3P and DHAP ^77^. Glucose metabolism is the major source of fuel in the brain ^37^. Through multiple steps, glucose is metabolized to pyruvate which is converted to acetyl-CoA to produce ATP through the TCA cycle ^37^. Our data suggest that alcohol reduces glycolysis and the TCA cycle in the NAc. These data are in line with the findings by Volkow and colleagues that glucose metabolism in the brain is significantly decreased in subjects suffering from AUD ^78–82^. In contrast to our results, mTORC1 was reported to increase glycolysis in skeletal muscle cells ^83^, macrophages ^84^, embryonic fibroblasts ^85^ and adipocytes ^86^. However, since mTORC1 is generally activated during periods of nutrient availability, we cannot exclude a potential feedback loop mechanism to constrain cellular energetic needs through mTORC1-dependent miR upregulation and Aldolase A repression. Our data showing that glycolysis is inhibited by alcohol via mTORC1, provide yet another unexpected role for mTORC1 and suggest again that mTORC1 function depends on context and/or cell type. In line with this possibility is our data showing that the activation of mTORC1 by alcohol in D1 NAc promotes the translation of microRNA machinery transcripts, resulting in the attenuation of Aldolase A transcription via miR-34a-5p which were also localized to D1 NAc neurons. Importantly, we found that miR34a-5p in D1 NAc neurons promotes alcohol but not sucrose intake. We cannot exclude the possibility that other targets of miR34a-5p in D1 NAc neurons contribute to alcohol drinking. However, putting together all our findings (**Fig. 9**), it is likely that the cell-specific attenuation of Aldolase A translation by miR34a-5p has a major contribution to neuroadaptations underlying alcohol drinking behavior. It will also be of interest to determine whether this cell-specific signaling pathway contributes to neuroadaptions that promote behaviors associated with other drugs of abuse.

Intriguingly, neuroadaptations that underlie addiction in general and AUD in specific ^87^ are high energy-consuming mechanisms that require ATP ^88^. The finding that glycolysis is inhibited by alcohol bears the question of what is the alternative energy source that is required for alcohol-dependent neuroadaptations in the NAc. Interestingly, mTORC1 was reported to increase the pentose phosphate pathway which could be an alternative energy source for neurons^89^. Another possibility is that acetyl-CoA, instead of being generated by pyruvate, is produced by medium-chain fatty acids (MCFAs) through fatty acid beta-oxidation ^90^. For instance, the medium-chain fatty acid octanoate can cross the blood-brain barrier ^91^, and has been shown to directly contribute ∼20% of energy in the rat brain ^92^. Thus, it is possible that when glucose levels are low, acetyl-CoA is produced through fatty acid metabolism. Another potential mechanism relates to acetate being metabolized from alcohol in the liver and astrocytes ^93–95^. Acetate can be converted to acetyl-CoA which enters the TCA cycle. Thus, acetate may be an alternative energy source in the NAc.

In addition to pyruvate being converted to acetyl-CoA, pyruvate is also the source of lactate the end product of glycolysis ^36,96^. We found that withdrawal from excessive alcohol intake reduces lactate content in the NAc, a process that requires mTORC1. This finding suggests a vectorial signaling in which alcohol withdrawal activates mTORC1 which increases the biogenesis of miR-34a-5p, in turn reducing the translation of Aldolase A leading to the attenuation of glycolysis and its final product, lactate (**Fig. 9**).

Lactate was initially thought to be produced solely by astrocytes and to be shuttled to neurons according to their energetic needs ^97^. However, the lactate shuttle model has been disputed in part due to discrepancies in stoichiometry and kinetics of lactate production ^98^. In fact, a large body of evidence have shown that glycolysis takes place in neurons ^37,88,99–103^, and we recently showed that hippocampal neurons metabolize glucose which is required for their normal function ^102^. As mTORC1 activation by alcohol occurs essentially in D1 neurons and that rapamycin administration effectively reverses the alcohol-mediated reduction in lactate levels, our model strongly suggests a specific reduction of lactate levels within D1 neurons.

For many years lactate was thought to be only a byproduct of glycolysis ^36^. However, recent studies showed that lactate is a signaling molecule and a substrate for epigenetic modification ^36,101,104^. For example, secreted lactate is a ligand for the Gi-coupled hydroxycarboxylic acid receptor 1 (HCAR1) ^105^, and lactate binding to HCAR1 modulates neuronal network function and synaptic plasticity ^105^. In addition, lactate was shown to be a substrate for lactylation, a newly identified posttranslational modification on proteins, including Histone H3 and Histone H1, resulting in enhanced gene transcription ^106–108^. Thus, it is plausible that alcohol-mediated reduction of lactylation is the reason for the large number of transcripts that are reduced by alcohol in the NAc.

Further studies reported that ketogenic diet which is given as a replacement for glucose reduces negative withdrawal symptoms in humans and mice ^109–111^ and alcohol intake in rats ^110^. Interestingly, ketogenic diet was shown to decrease mTORC1 activity in the hippocampus and the liver ^112,113^. These data, together with ours, suggest an intriguing possibility that, like the hypodopaminergic state in AUD subjects, which drives further drinking to alleviate allostasis symptoms, hypoglycolysis also promotes further drinking to supply the brain with an alternative energy source. Further investigation on this topic is warranted.

Finally, we show that subcutaneous administration of L-lactate significantly reduces alcohol consumption. Attenuation of lactate levels or its shuttling between astrocytes and neurons is associated with stress ^107^ and depression ^114,115^. Importantly, L-lactate administration reduces depressive-like symptoms in mice ^116^, and we found that systemic administration of L-lactate in mice reduces alcohol but not sucrose intake. Together, these data and ours suggest that psychiatric disorders are associated with an imbalance in lactate levels in the brain and that adjusting lactate levels reduces adverse effects associated with alcohol and depression. Furthermore, these data provide preclinical data to suggest that L-lactate could be developed as a cost-effective readily available approach to treat AUD and potentially other psychiatric disorders.

In summary, this study unveils a novel dimension of mTORC1 function in alcohol-related behaviors. Specifically, our study unravels a paradoxical action of mTORC1 in orchestrating both translation repression and metabolic shift in response to chronic alcohol exposure which in turn drives further alcohol intake.

## Methods

### Animals and Breeding

Male C57BL/6J mice (6-8 weeks old at time of purchase) were obtained from The Jackson Laboratory. Drd1a-Cre (D1-Cre) and AdoraA2-Cre (A2A-Cre) mice both of which are on C57BL/6 background, were obtained from Mutant Mice Resource and Research Centers (MMRRC) UC Davis. Ribotag mice (ROSA26CAGGFP-L10a), which express the ribosomal subunit RPL10a fused to EGFP (EGFP-L10a) in Cre-expressing cells ^117^, were purchased from The Jackson Laboratory (B6;129S4-Gt (ROSA)26Sortm9(EGFP/Rpl10a)Amc/J). Ribotag mice were crossed with D1*-*Cre mice allowing EGFP-L10a expression in D1*-*expressing cells. Mouse genotype was determined by poly-chain reaction (PCR) analysis of tail DNA.

Only males (age 6-8 weeks) were used in the study. Mice were individually housed on paper-chip bedding (Teklad #7084), under a reverse 12-hour light-dark cycle (lights on 1000 to 2200 h). Temperature and humidity were kept constant at 22 ± 2°C, and relative humidity was maintained at 50 ± 5%. Mice were allowed access to food (Teklad Global Diet #2918) and tap water *ad libitum*. All animal procedures were approved by the university’s Institutional Animal Care and Use Committee and were conducted in agreement with the Association for Assessment and Accreditation of Laboratory Animal Care.

### Plasmids generation viral production

The mouse miR-34a nucleotide sequence (5’CCAGCTGTGAGTAATTCTTTGGCAGTGTCTTAGCTGGTTGTTGTGAGTATTAGCTAAGGAAGCAATCAGCAAGTATACTGCCCTAGAAGTGCTGCACATTGT3’) was synthesized. Synthesized DNA oligos containing the miRNA sequences were annealed and inserted into pLL3.7 vector (11795, Addgene) at HpaI and XhoI sites. Plasmids were prepared using a Plasmid Maxi Kit. All constructs were verified by sequencing.

The production of miR-34a5p lentivirus was conducted as described in ^19^. Briefly, HEK293 lentiX cells (Clontech, Mountain View, CA) were transfected with the lentiviral packaging vectors psPAX2 and pMD2.G, together with the pLL3.7 miR-34a-5p or pLL3.7 GFP using lipofectamine 2000 in Opti-MEM medium. Six hours after transfection, medium was replaced to DMEM-FBS 10%. Sixty hours after transfection, supernatant containing the viral particles was collected, filtered into 0.22µm filters and purified by ultracentrifugation at 26.000 g for 90 minutes at 4°C. The pellet fraction containing the virus was resuspended in sterile PBS, aliquoted and stored at −80°C until use. Virus titer was determined using the HIV-1 p24 antigen ELISA kit.

The plasmid and Adeno-associated virus (AAV)2-CMV-Flex-miR-34a-EGFP-WPRE (AAV2-Flex-miR-34a, titer: 2.89×10^13^ GC/ml) were designed and produced by the VectorBuilder.

### Tissue harvesting

Mice were killed and brains were rapidly removed on an anodized aluminum block on ice. The NAc was isolated from a 1 mm thick coronal section located between +1.7 mm and +0.7 mm anterior to bregma according to the Franklin and Paxinos stereotaxic atlas (3rd edition). Collected tissues were immediately homogenized in 300 µl radioimmuno-precipitation assay (RIPA) buffer (50 mM Tris-HCl, pH 7.6, 150mM NaCl, 2mM EDTA, 1% NP-40, 0.1% SDS. and 0.5% sodium deoxycholate and protease and phosphatase inhibitors cocktail). Samples were homogenized by a sonic dismembrator. Protein content was determined using a BCA kit.

### Polysomal fractionation

Polysome-bound RNA was purified from mouse NAc according to a protocol we described previously ^18^. Specifically, fresh mouse NAc was snap-frozen in a 1.5ml Eppendorf tube and pulverized in liquid nitrogen with a pestle. After keeping on dry ice for 5 minutes, the powder of one NAc was resuspended in 1 ml lysis buffer (10mM Tris pH 8.0, 150mM NaCl, 5mM MgCl_2_, 1% NP40, 0.5% sodium deoxycholate, 40mM dithiothreitol, 400U/ml Rnasin, 10mM Ribonucleoside Vanadyl Complex and 200µg/ml cycloheximide) followed by pipetting 20 times to further disrupt cell membranes. The homogenate was centrifuged for 10 seconds at 12.000g to remove intact nuclei. The supernatant was collected, and ribosomes were further released by adding 2X extraction buffer (200mM Tris pH7.5, 300mM NaCl and 200µg/ml cycloheximide). Samples were kept on ice for 5 minutes and then centrifuged at 12.000g, 4°C for 5 minutes to remove mitochondria and membranous debris. The resulting supernatant was loaded onto a 15%-45% sucrose gradient and centrifuged in a SW41Ti rotor at 38.000rpm, 4°C for 2 hours. Sucrose gradient fractions were collected and further digested with proteinase K solution (400µg/ml proteinase K, 10mM EDTA, 1% SDS) at 37°C for 30 minutes, followed by phenol-chloroform extraction. RNA in the water phase of the polysomal fraction was recovered by ethyl alcohol precipitation. The integrity of the polysomal fractions is described in ^18^. Specifically, RNA was visualized by migrating on a 1.5% agarose gel. Fractions 01-05 mainly contain tRNAs. Fractions 06-15 are enriched with 40S ribosomal subunit as well as 18S rRNA. Fractions 16-25 are enriched with 60S ribosomal subunit and 28S rRNA. Fractions 26-40 contain polysomal RNA^18,20^.

### cDNA synthesis and real-time quantitative PCR (RT-qPCR)

Total RNA extracted from tissues was treated with DNase I. Synthesis of cDNA was performed using the iScript cDNA Synthesis Kit according to the manufacturer’s instructions (Biorad). For polysomal RNA, and TRAP D1 RNA, cDNA synthesis and amplification were conducted using Ovation RNA Amplification Kit V2 (Tecan). The resulting cDNA was used for quantitative RT-qPCR, using SYBR Green PCR Master mix (Thermo Fisher Scientific). Thermal cycling was performed on QuantStudio 5 real-time PCR System using a relative calibration curve. The quantity of each mRNA transcript was measured and expressed relative to Glyceraldehyde-3-Phosphate dehydrogenase (GAPDH). The Primers are listed in **Supplementary Table 4**.

### *In Silico* miR prediction

miRNA predictions were based on TargetScan 2 ^118^, miRwalk ^119^ and miRDB ^120,121^. miR-34a-5p and Aldolase A complimentary binding site was predicted using miRWalk ^119^. miRWalk uses TarPmiR, a predictive algorithm, to generate a binding score ^119^. Specifically, TarPmiR applies the trained random forest-based predictor to determine the target sites ^122^.

### miRNA extraction, cDNA synthesis and quantitative RT-qPCR

microRNAs were extracted from tissues using miRNeasy Mini Kit according to the manufacturer’s instructions (Qiagen). miRNAs yield and purity was evaluated using a nanodrop ND-1000 spectrophotometer. cDNA synthesis was performed using the miRCURY LNA RT kit according to the manufacturer’s instructions (Qiagen), starting with 500ng of RNA and 5X reaction mix. Enzyme and nuclease-free water were added to a final volume of 20µl. RNA spike-in of Unisp6 was added to each sample to monitor the efficiency of the reverse transcription reaction. Quantitative RT-qPCR was performed using miRCURY LNA SYBR Green master mix according to the manufacturer’s instructions (Qiagen), and PCR samples were run on the QuantStudio 5. Each data point represents an average of 3 replicates. Relative microRNA expression was determined by the following calculation according to ^123^. First, the average of cycle threshold (CT) of the miRNA of interest was subtracted from the CT of the control U6 rRNA to generate ΔCT. ΔCTs were then subtracted from the average of control ΔCT values for each sample to generate the ΔΔCT. miR levels were then calculated as 100 multiplied by 2^-ΔΔCT^ ± S.E.M. The Primers are listed in **Supplementary Table 4**.

### TRAP purification

After NAc dissection, D1 **or D2-**specific mRNAs were purified according to the established TRAP protocol ^124^. The NAc was homogenized with a glass homogenizer in an ice-cold lysis buffer (150mM KCl, 20mM HEPES [pH 7.4], 10mM MgCl2, 0.5mM dithiothreitol, 100µg/mL cycloheximide, 80U/µl RNasin Plus Rnase Inhibitor, and EDTA-free protease inhibitors). Following homogenization, samples were centrifuged at 2,000g at 4°C for 10 minutes and the supernatant was removed to a new tube. NP-40 (final concentration 1%) and 1,2-Diheptanoylsn-glycero-3-phosphocholine (DHPC, final concentration 15 mM) were subsequently added and samples were incubated on ice for 5 minutes. Samples were centrifuged at 20,000g at 4°C for 10 minutes and the supernatant was transferred to a new tube. Streptavidin Dynabeads coated with biotin-linked mouse anti-GFP antibodies were then added to the supernatant and the samples were incubated overnight at 4°C with end-over-end rotation. Beads were collected on a magnetic rack and washed three times with wash buffer (350mM KCl, 20 mM HEPES pH 7.4, 10 mM MgCl_2_, 0.5 mM dithiothreitol, 100µg/mL cycloheximide, 1% NP-40). RNA was subsequently purified using the Absolutely RNA Isolation Nanoprep kit (Agilent). To ensure accurate quantitation, purified RNA was run on a Qubit 4 (Thermofisher). The validation of the RiboTag specificity is shown in **Supplementary Fig. 13**.

### Western blot analysis

Equal amounts of homogenates from individual mice (30µg) were resolved on NuPAGE Bis-Tris gels (4%-12% gradient, Life Technologies) and transferred onto nitrocellulose membranes (Millipore). Blots were blocked in 5% milk-PBS and 0.1% Tween 20 for 30 minutes and then incubated overnight at 4°C with primary antibodies. Membranes were then washed and incubated with HRP-conjugated secondary antibodies for 2 hours at room temperature. Bands were visualized using Enhanced Chemiluminescence (ECL, Millipore). The optical density of the relevant band was quantified using ImageJ 1.44c software (NIH). Antibodies details are listed in **Supplementary Table 5**.

### Immunochemistry

Mice were deeply anesthetized with Euthasol and perfused with 0.9% NaCl, followed by 4% paraformaldehyde in PBS, pH 7.4. Brains were removed, post-fixed in the same fixative for 2 hours, and transferred to PBS at 4°C. On the following day, brains were transferred into 30% sucrose and stored for 3 days at 4°C. Thirty µm-thick coronal sections were cut on a cryostat, collected serially and stored at −80°C. Sections were permeabilized with, and blocked in, PBS containing 0.3% Triton and 5% donkey serum for 4 hours. Sections were then incubated for 18 hours at 4°C on an orbital shaker with anti-pS6 (1:500) and anti-NeuN antibodies (1:500) diluted in 3% bovine serum albumin (BSA) in PBS. Next, sections were washed in PBS then incubated for 4 hours with Alexa Fluor 596-labeled donkey anti-rabbit and Alexa Fluor 647-labeled donkey anti-mouse diluted in 3% BSA in PBS. After staining, sections were rinsed in PBS and cover slipped using Prolong Gold mounting medium (Thermofisher). Images were acquired using an Olympus Fluoview 3000 Confocal microscope using manufacture recommended filter configurations. Quantification was performed using the cell counter plugin in ImageJ software (NIH). Antibodies details are listed in **Supplementary Table 5**.

### Luciferase assay

The 3′ untranslated region (UTR) of Aldolase A containing miR-34a-5p predicted target site was cloned into the pmirGLO Dual-Luciferase miRNA Target Expression Vector (E1330, Promega). Mutant construct was generated with a mutated target site. HEK293 cells were seeded in 96-well plates and co-transfected with luciferase reporters, mimic miR-34a-5p (4464066, Invitrogen), or a mimic miR negative control (4464058, Invitrogen). Firefly (FL) and Renilla (RL) luciferase activities were measured 48 hours after transfection using TECAN plate reader. Signal was calculated using FL/RL ratio relative to the empty pmirGLO reporter vector.

### NAc shell viral infection

Intra-NAc shell infusion of lentivirus or AVV2 was conducted as described in ^19^. Briefly, mice were anesthetized using isoflurane. Bilateral viral infusions were done using stainless steel injectors (33 gauge, Hamilton) into the NAc shell (anteroposterior +1.2 mm, mediolateral ± 0.75 mm and dorsoventral −4.30 mm, from bregma). Animals were infused with either Ltv-Control expressing GFP only or Ltv-miR-34a-5p (1.2×10^8^ pg/ml, 1µl/side), or **AAV2-Flex-control or AAV2-Flex-miR-34a (2.89×10**^13^ **GC/ml, 1µl/side)** at an infusion rate of 0.1µl/minute. After each infusion, the injectors were left in place for an additional 10 minutes to allow the virus to diffuse.

### Preparation of solutions

Alcohol solution was prepared from absolute anhydrous alcohol (190 proof) diluted to 20% alcohol (v/v) in tap water. Sucrose solution (1%) was dissolved in tap water (w/v).

Rapamycin (20mg/kg, R-5000, LC Laboratories) was dissolved in 5% DMSO and 95% saline. Vehicle contained 5% DMSO and 95% saline. Sodium L-Lactate (2g/kg, Sigma Aldrich, 867-56-1) and NaCl (1g/kg) were dissolved in PBS as described in ^41^.

### Drug administration

Rapamycin: rapamycin (20mg/kg) or vehicle was administered i.p. 3 hours before the end of the last alcohol withdrawal session and tissues were harvested at the end of the 24 hour withdrawal session as in ^19^.

L-lactate: L-lactate (2g/kg) or NaCl (1g/kg) was administered subcutaneously (s.c.). A “within-subject” design in which mice received both treatments in counterbalanced order, with one week in between treatments. Specifically, on weeks 8 and 9, mice were administered s.c. with L-lactate (2g/kg), or vehicle solution 30 minutes before the beginning of the drinking session. On weeks 10 and 11, mice were systemically administered with NaCl (1g/kg), or vehicle 30 minutes before the beginning of the drinking session. Alcohol and water consumption was evaluated at the end of 4 hours and 24 hours drinking session.

### Lactate measurement

The NAc lysates were added to 96-well plates and adjusted to 50µl of reaction mix as described in the lactate colorimetric assay kit instructions (L-Lactate Assay Kit ab65331, Abcam). After incubation for 30 minutes at room temperature in the dark, the absorbance at 570 nm was measured using a microplate reader.

### Metabolomics

Metabolomics analysis was conducted as described in ^102^. Specifically, fresh mouse NAc was snap-frozen in a 1.5ml Eppendorf tube in liquid nitrogen. To extract the metabolites from the frozen tissue, samples were first homogenized in a cryogenic mortar and pestle before being mixed with 1ml of 80% methanol chilled to −80°C. Samples were then vortexed for 20 seconds and incubated at −80°C for 20 minutes. Following the incubation, samples were vortexed for an additional 20 seconds and then centrifuged at 16,000g for 15 minutes at 4°C. The supernatant was transferred to a −80°C prechilled tube. BCA assay was used to normalize the extracted metabolites to protein content. A 100µg protein equivalent of extracted metabolites was aliquoted from each sample and dried in a CentriVap. The dried samples were then stored at - 80°C until analysis by the UCLA Metabolomics Center.

### Drinking paradigm

#### Two bottle choice - 20% alcohol

Mice underwent 7 weeks of intermittent access to 20% (v/v) alcohol in a two-bottle choice drinking paradigm (IA20%2BC) as previously described ^22^. Specifically, mice had 24-hour access to one bottle of 20% alcohol and one bottle of water on Mondays, Wednesdays, and Fridays, with alcohol drinking sessions starting 2 hours into the dark cycle. During the 24 or 48 hours (weekend) of alcohol withdrawal periods, mice had access to a bottle of water. The placement (right or left) of the bottles was alternated in each session to control for side preference. Two bottles containing water and alcohol in an empty cage were used to evaluate the spillage. Alcohol and water intake were measured at the end of each 24 hours drinking session (**Supplementary Tables 1**).

#### Two bottle choice - 1% sucrose

Mice underwent 2 weeks of intermittent access to 1% (v/v) sucrose in a two-bottle choice drinking paradigm as previously described ^19^. Specifically, mice had 24-hour access to one bottle of 1% sucrose and one bottle of water on Mondays, Wednesdays, and Fridays, with sucrose drinking sessions starting 2 hours into the dark cycle. The placement (right or left) of the bottles was alternated in each session to control for side preference. Two bottles containing water and sucrose in an empty cage were used to evaluate the spillage. Sucrose and water intake were measured at the end of each 24-hour drinking session.

### Open field locomotion

Locomotion test was performed as described in ^125^. Specifically, mice were habituated for 5 minutes in the open field apparatus (43 x 43 cm). Mice were then injected s.c. with 2g/kg L-lactate or saline before being placed back in the open field apparatus and their movement was recorded for an additional 20 minutes. Mice were automatically video tracked using Ethovision software and locomotion per 1-minute bins was calculated.

### Glucose tolerance test

Glucose tolerance assay was performed as described previously ^22^. Briefly, mice were deprived of food for 6 hours and were then injected i.p. with 1 g/kg glucose. Blood samples were taken from a tail vein nick at different time intervals (0, 15, 30, 60, and 120 minutes post glucose administration), and blood glucose level was analyzed using a Bayer Contour blood glucose meter and test strips.

### Statistical analysis

GraphPad Prism 7.0 (GraphPad Software Inc., La Jolla, CA) was used to plot and analyze the data. D’Agostino–Pearson normality and F-test/Levene tests were used to verify the normal distribution of variables and the homogeneity of variance, respectively. Data were analyzed using the appropriate statistical test, including two-tailed paired t-test, two-tailed unpaired t-test, one-way analysis of variance (ANOVA), and two-way ANOVA followed by post hoc tests as detailed in **Supplementary Table 2**. All data are expressed as mean ± SEM, and statistical significance was set at p < 0.05.

## Data availability

Any data not contained in the paper are available on request with appropriate agreements. Source data are provided with this paper.

## Acknowledgments

This study was supported by the National Institute of Alcohol Abuse and Alcoholism, R01 AA027474 (D.R.), RF1 AG064170 and R01 AG065428 (K.N.) and F32 AG082460 (Y.J.S.). We thank Chhavi Shukla (University of California San Francisco) for technical assistance.

---

**Supplementary Figure 1.**
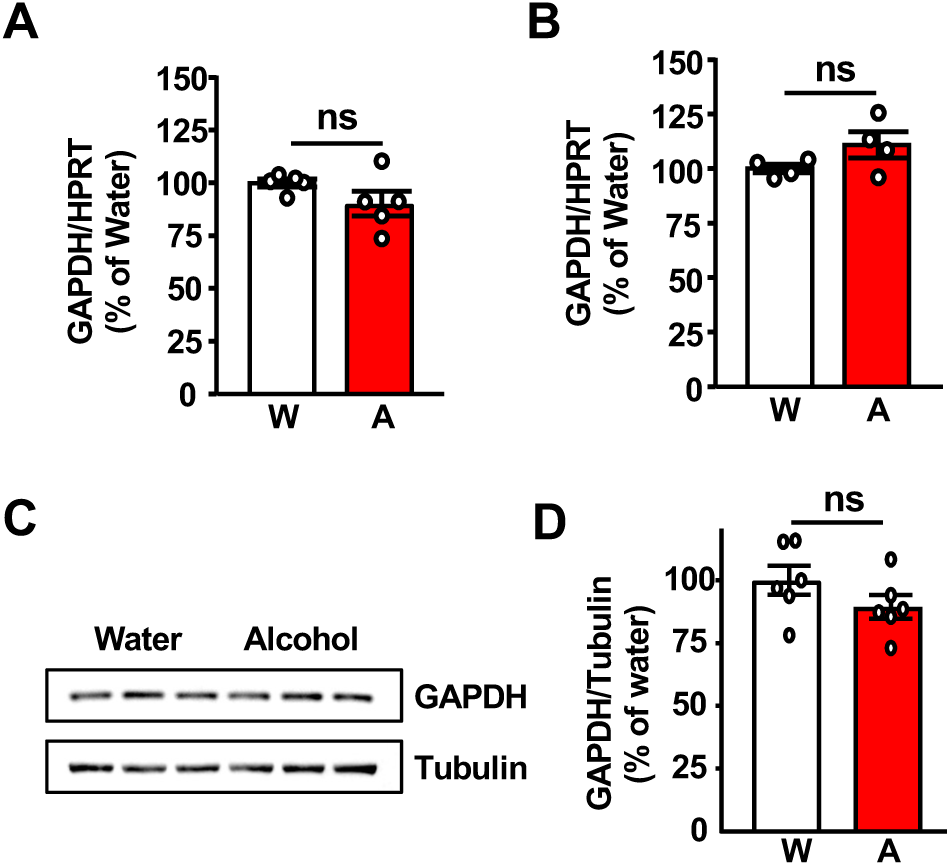
Alcohol does not alter GAPDH mRNA levels in the NAc. (**A-B**) Polysomal mRNA levels of GAPDH in D1+ neurons (**A**), and D2+ neurons (**B**) after alcohol withdrawal were determined by RT-qPCR. Data are presented as the average ratio of a transcript to HPRT±SEM and expressed as % of water control. Statistical analyses are depicted in **Supplementary Table 2**. ns: non-significant. (**C**) GAPDH protein levels in the NAc. (**D**) Data are presented as the average ratio of GAPDH to Tubulin±SEM and are expressed as % of water control. Statistical analyses are depicted in **Supplementary Table 2**. ns: non-significant.

**Supplementary Figure 2.**
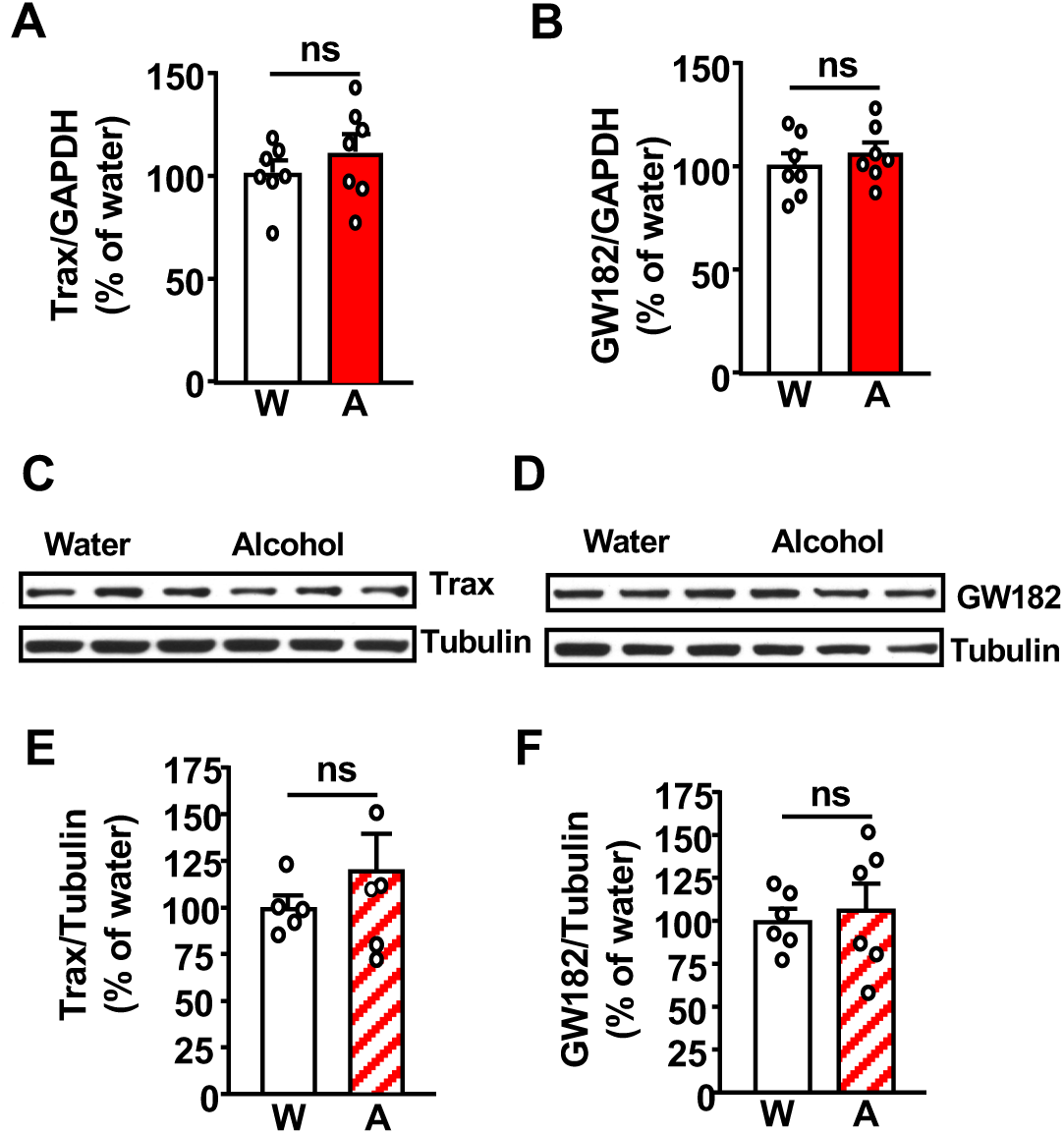
Alcohol does not alter the transcription of Trax and GW182 in the NAc and the protein levels in the DLS. After 7 weeks of IA20%2BC (**Supplementary Table 1**), the NAc of mice was dissected at the end of the 24 hours alcohol withdrawal session. Control animals had access to water only. **(A-B)** GW182 and Trax mRNA level was determined by RT-qPCR in the total mRNA sample. Data are presented as the average ratio of GW182 or Trax to GAPDH±SEM and expressed as % of water control. Statistical analyses are depicted in **Supplementary Table 2**. ns: non-significant. **(C-D)** Trax and GW182 protein levels in the DLS. (**E-F**) Data are presented as the average ratio of GW182 or Trax to Tubulin±SEM and are expressed as % of water control. Statistical analyses are depicted in **Supplementary Table 2**. ns: non-significant.

**Supplementary Figure 3.**
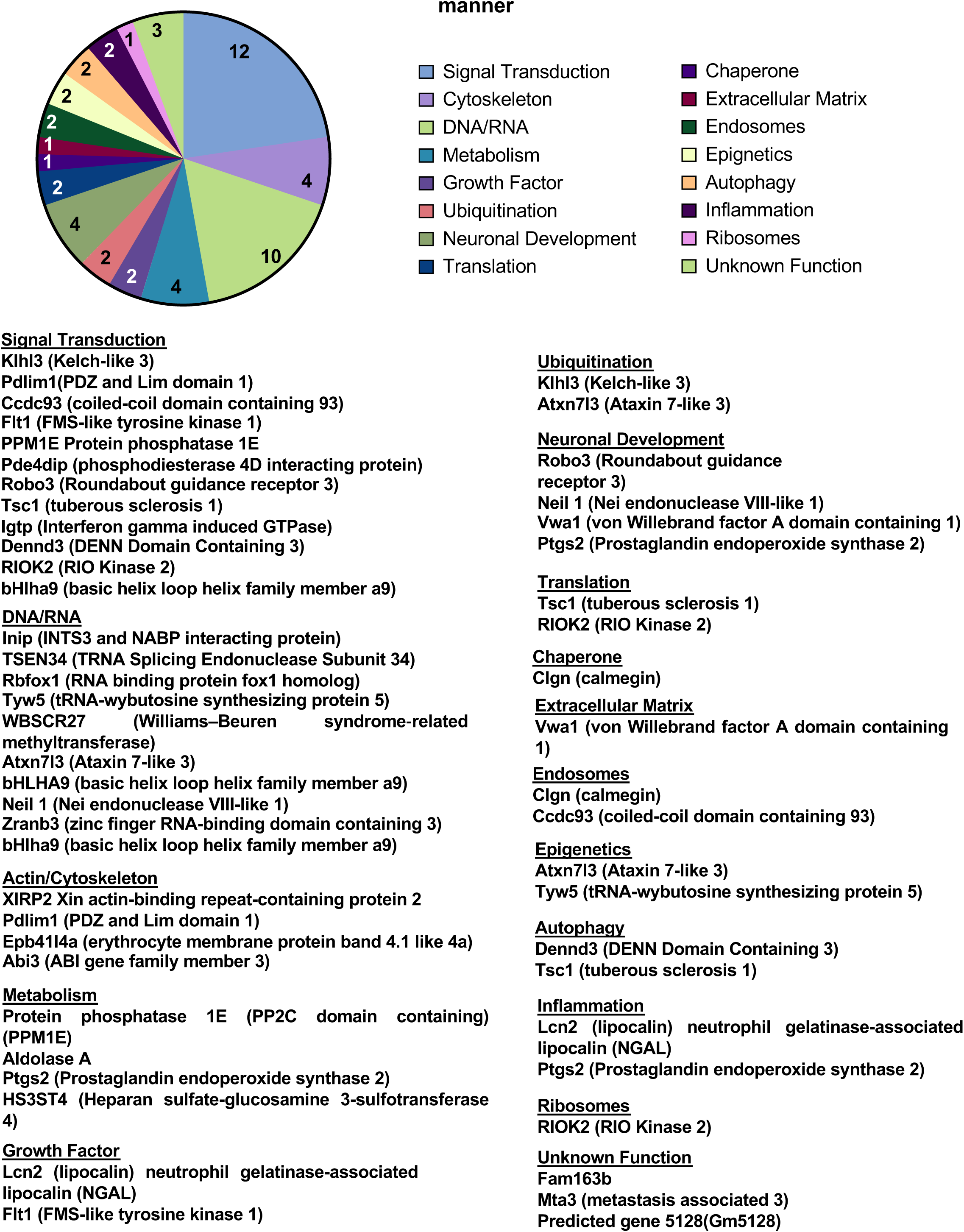
Functional characterization of RNAseq transcripts whose translation are decreased by alcohol in an mTORC1-dependent manner. Functional categories were determined by an up-to-date literature search of each of the transcripts highlighting the transcripts’ known function. Some proteins have several functions and are therefore found in multiple functional categories.

**Supplementary Figure 4.**
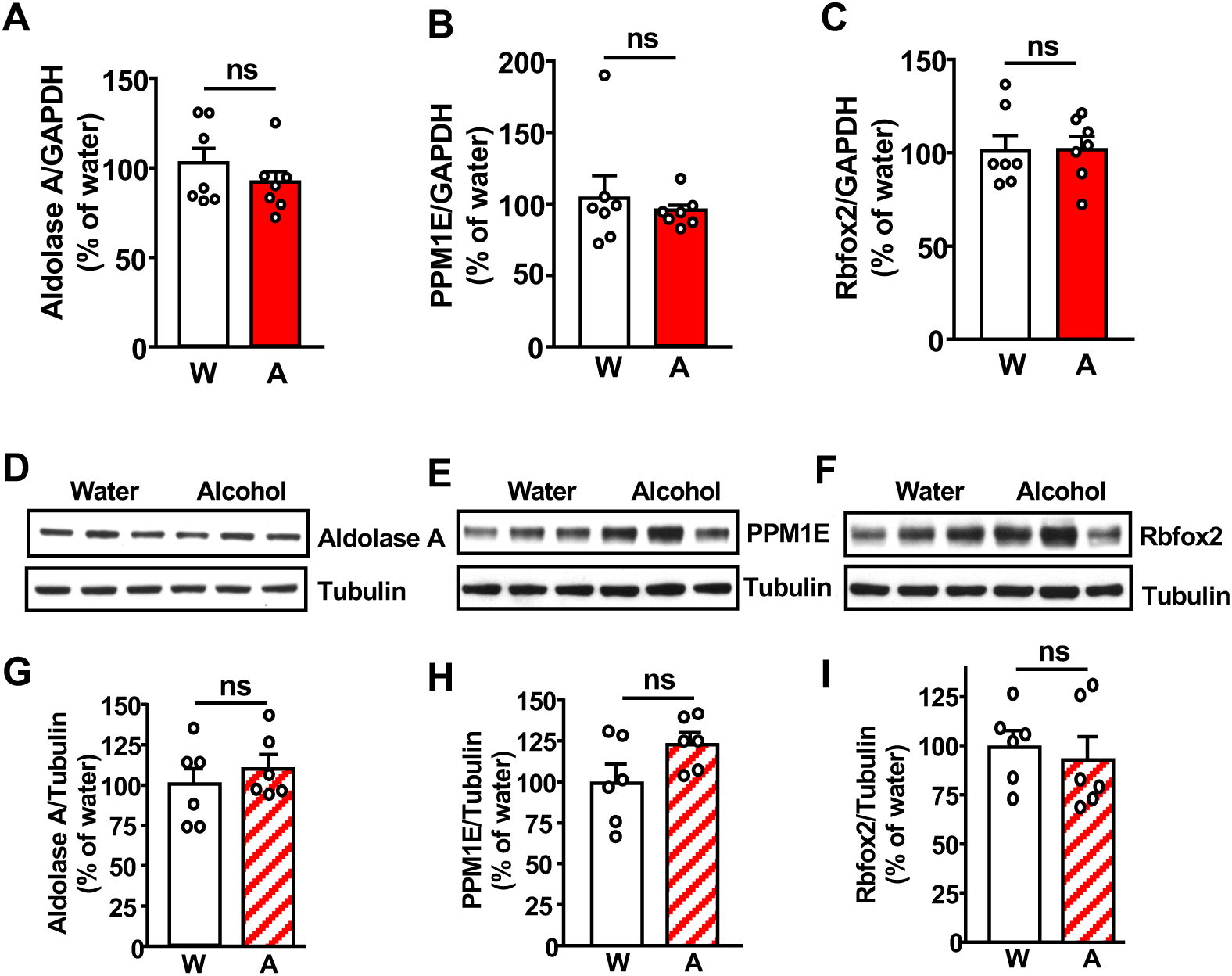
Alcohol does not alter the transcription of Aldolase A, PPM1E, Rbfox2 in the NAc and their protein level in the DLS. After 7 weeks of IA20%2BC (**Supplementary Table 1**), the NAc of mice was dissected at the end of the 24 hours alcohol withdrawal session. Control animals had access to water only. **(A-C)** Total Aldolase A (**A**), PPM1E (**B**), Rbfox2 (**C**) mRNA levels were determined by RT-qPCR in the total mRNA sample. Data are presented as the average ratio of Aldolase A, Rbfox2 or PPM1E to GAPDH±SEM and expressed as % of water control. Statistical analyses are depicted in **Supplementary Table 2**. ns: non-significance. **(D-F)** Aldolase A, PPM1E, Rbfox2 protein levels after withdrawal were determined by western blot analysis. (**G-I**) Data are presented as the average ratio of Aldolase A, PPM1E, Rbfox2 to Tubulin±SEM and are expressed as the % of water control. Statistical analyses are depicted in **Supplementary Table 2**. ns: non-significant.

**Supplementary Figure 5.**
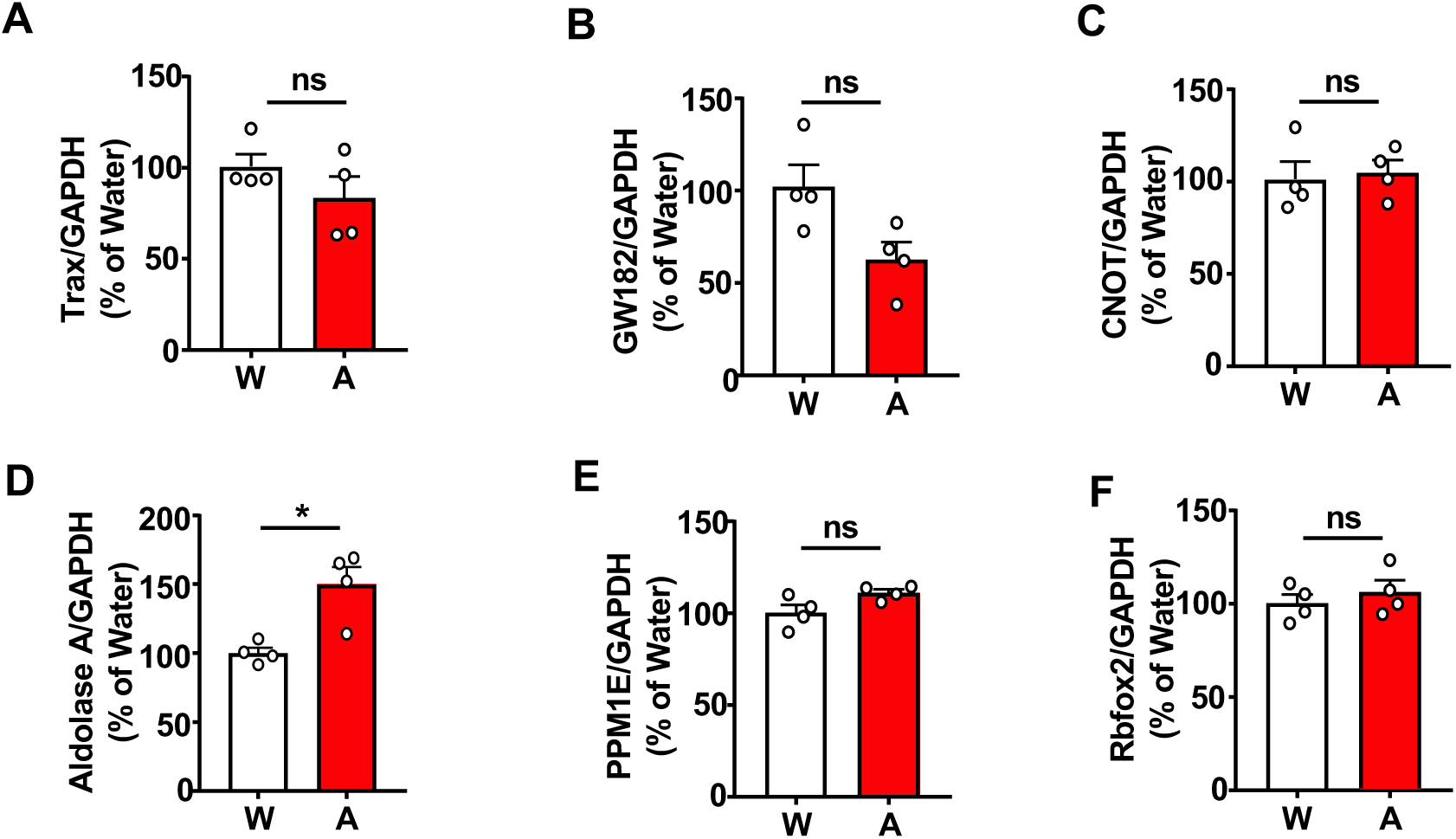
Alcohol does not affect the translation of Trax, Gw182 and CNOT4, PPM1E and Rbfox2 but increases the translation of Aldolase A in D2+ neurons. (**A-F**) Polysomal mRNA levels in D2+ neurons of Trax (**A**), GW182 (**B**), CNOT4 (**C**), Aldolase A (**D**), PPM1E (**E**), Rbfox2 (**F**) after alcohol withdrawal were determined by RT-qPCR. Data are presented as the average ratio of a transcript to GAPDH±SEM and expressed as % of water control. Statistical analyses are depicted in **Supplementary Table 2**. *p<0.05, ns: non-significant.

**Supplementary Figure 6.**
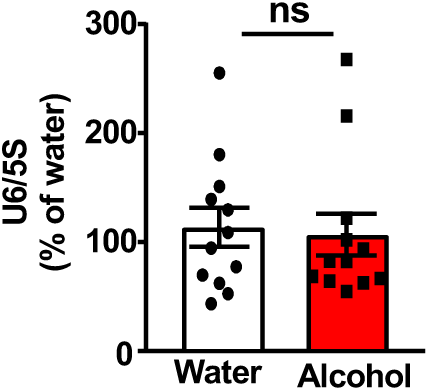
U6 expression in the NAc is not altered by alcohol. After 7 weeks of IA20%2BC (**Supplementary Table 1**), the NAc of mice was dissected at the end of the 24 hours alcohol withdrawal session. Control animals had access to water only. U6 levels were measured by RT-qPCR. Data are presented as the average ratio of U6 to 5S±SEM and expressed as % of water control. Statistical analyses are depicted in **Supplementary Table 2**. ns: non-significant.

**Supplementary Figure 7.**
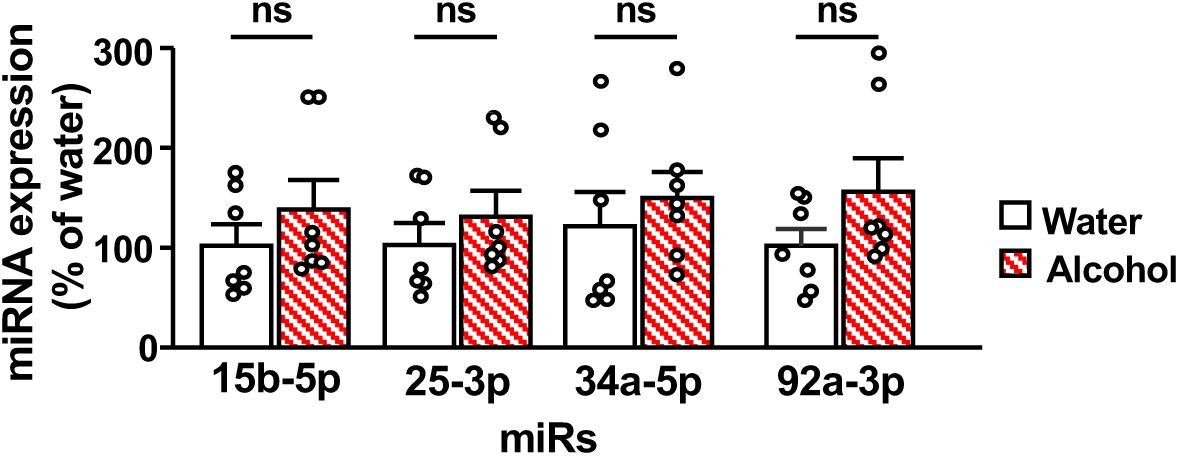
The levels of miR-15b-5p, miR-25-3p, miR-92a-3p, and miR-34a-5p are not altered in the DLS. After 7 weeks of IA20%2BC (Supplementary Table 1), the NAc of mice was dissected at the end of the 24 hours alcohol withdrawal session. Control animals had access to water only. miRs levels were measured by RT-qPCR. Statistical analyses are depicted in Supplementary Table 2. ns: non-significant.

**Supplementary Figure 8.**
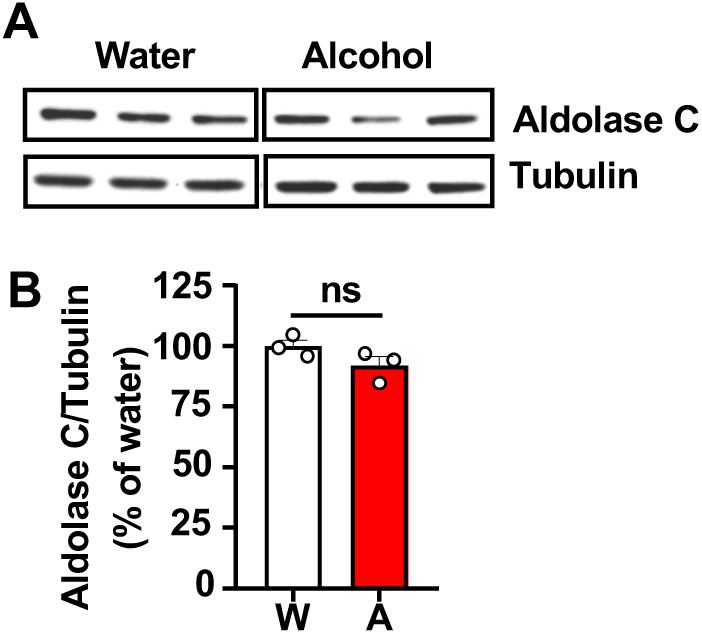
Alcohol does not alter the protein levels of Aldolase C in the NAc. After 7 weeks of IA20%2BC (**Supplementary Table 1**), the NAc of mice was dissected at the end of the 24 hours alcohol withdrawal session (A, red). Control animals had access to water only (W, white). **(A)** Aldolase C level was determined by western blot analysis. (**B**) Data are presented as the average ratio of Aldolase C to Tubulin±SEM and are expressed as % of water control. Statistical analyses are depicted in **Supplementary Table 2**. ns: non-significant.

**Supplementary Figure 9.**
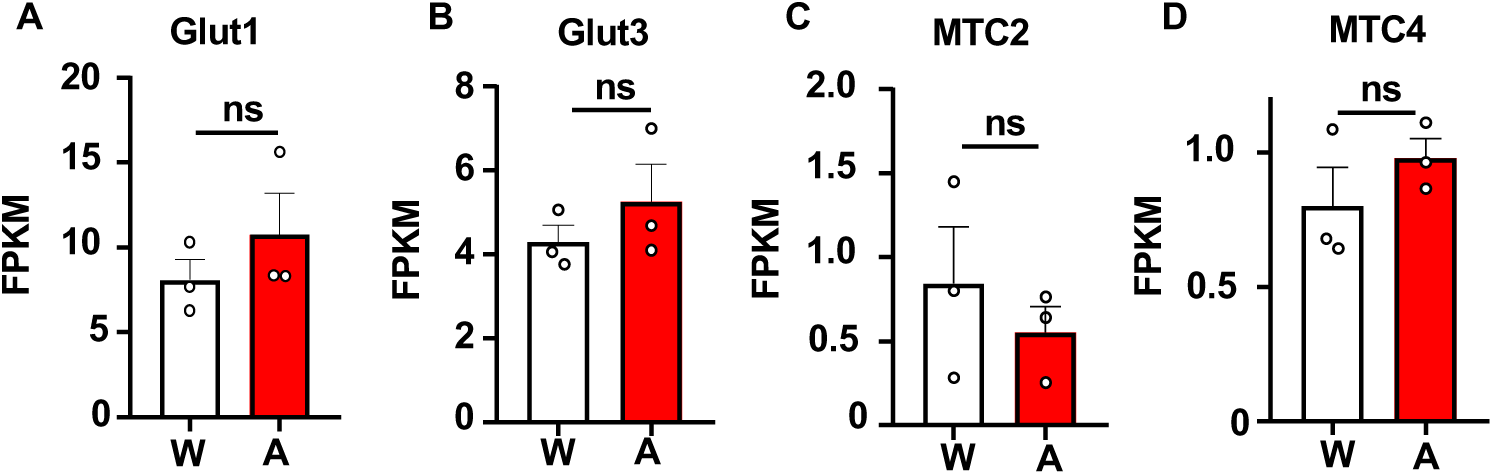
Alcohol does not alter the translation of the main glucose and lactate transporters in the NAc. After 7 weeks of IA20%2BC, the NAc of mice was dissected at the end of the 24 hours alcohol withdrawal session (A, red). Control animals had access to water only (W, white). RNAseq data ^19^ revealed that the translation of the astrocytic glucose transporter Glu1 (**A**), the neuronal glucose transporter Glu3 (**B**), the neuronal lactate transporter MTC2 (**C)** and the astrocytic lactate transporter MTC4 **(D**) is unaltered by alcohol. W: water, A: alcohol, n=3. ns: non-significant. FPKM: Fragments per kilobase of transcript per million mapped reads

**Supplementary Figure 10.**
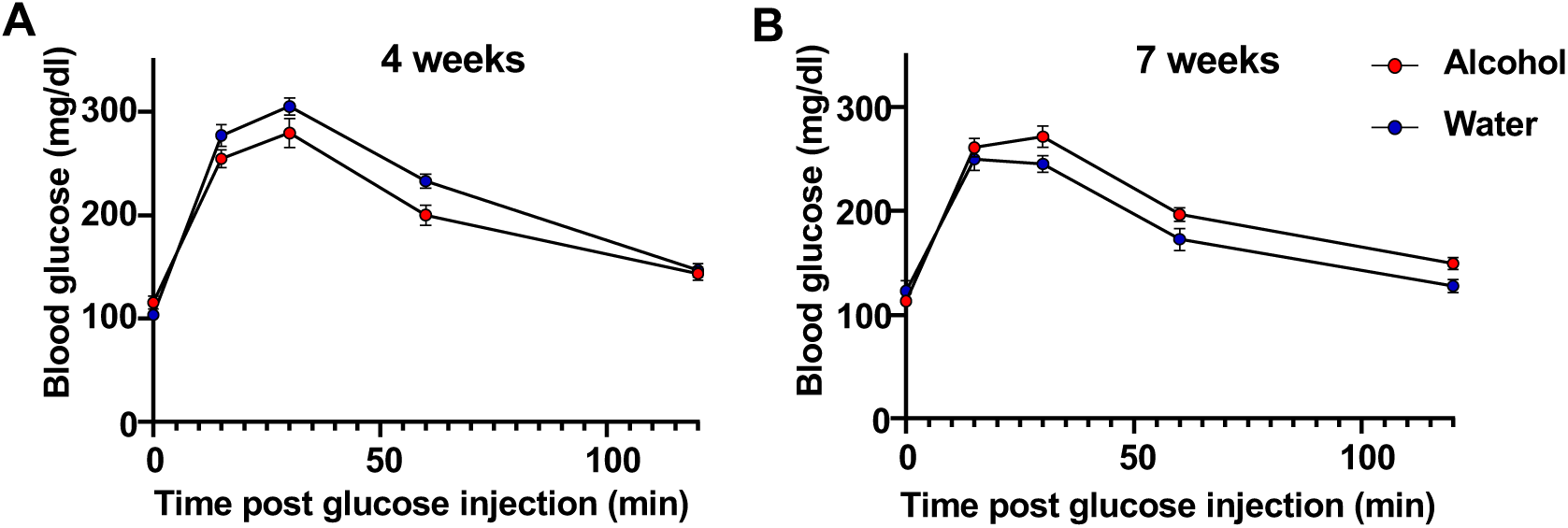
Blood glucose tolerance is unaltered by chronic alcohol drinking. (**A-B**) Glucose tolerance test was performed after 4 weeks (**A**) and 7 weeks (**B**) of IA20%2BC (**Supplementary Table 1**). Control animals had access to water only. Mice were subjected to 6 hours fasting before glucose (1g/kg) injection. Blood was collected before glucose injection 15-120 minutes post glucose administration, and glucose levels were determined. Statistical analyses are depicted in **Supplementary Table 2**.

**Supplementary Figure 11.**
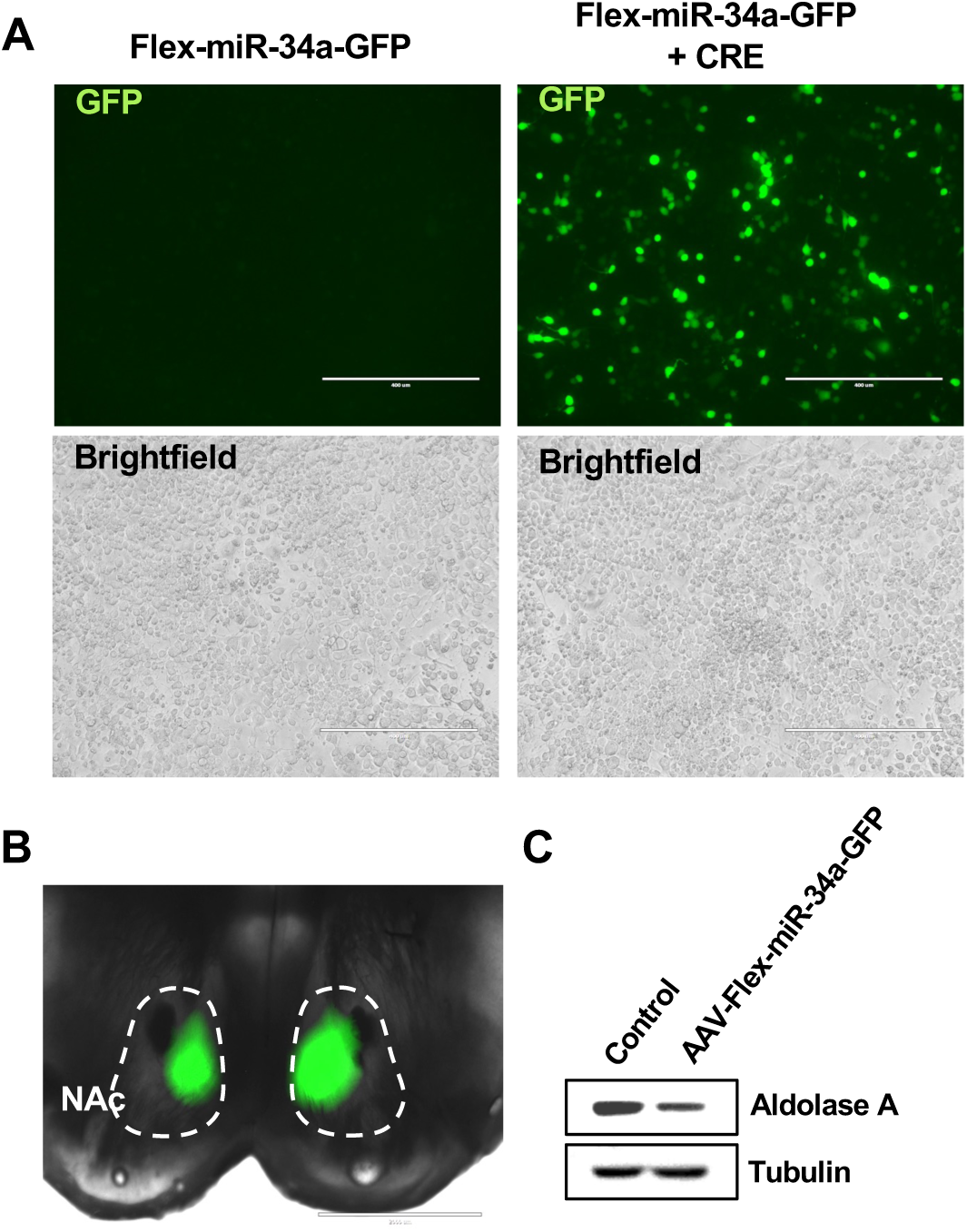
Characterization of AAV2-Flex-miR-34a-GFP in cells and in the NAc. **(A)** N2A cells were transfected with plasmid pAAV2-Flex-miR-34a-GFP alone or combined with the plasmid pAAV-Cre. Representative images at 20x magnification of labeled GFP in green, and brightfield. Bar scale 400µm. **(B-C)** AAV-Cre-dependent overexpression of miR34a in D1 NAc neurons. AAV2-Flex-miR-34a-GFP or AAV2-Flex-control was infused into the NAc of D1-Cre mice and Aldolase A protein level was evaluated by western blot analysis.

**Supplementary Figure 12.**
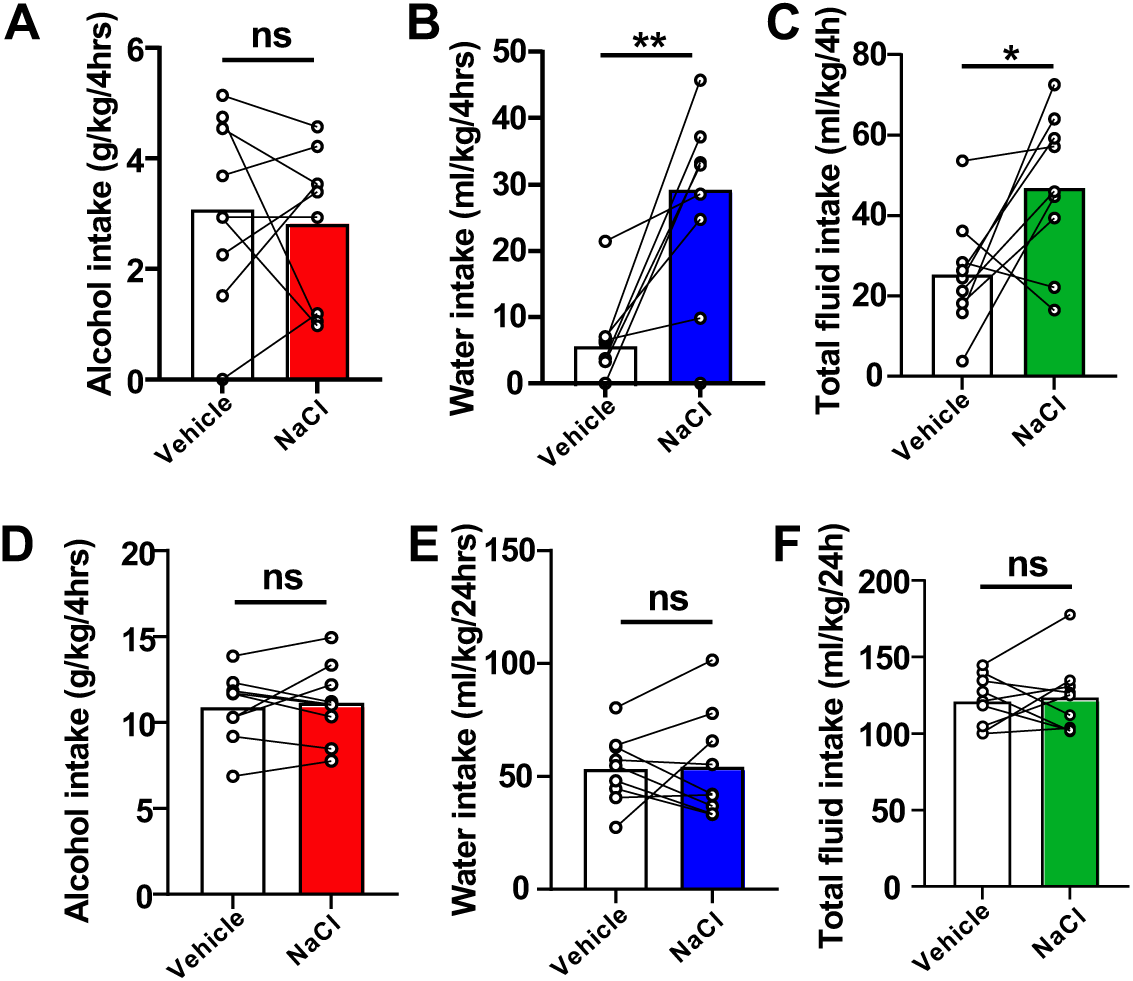
Subcutaneous administration of NaCl does not affect alcohol consumption. Mice underwent 7 weeks of IA20%2BC (**Supplementary Table 1**) whereas control animals had access to water bottles only. On weeks 10 and 11, mice were s.c. injected with NaCl (1g/kg) or PBS in a counterbalanced manner 30 minutes before the beginning of a 24-hour drinking session. Alcohol and water consumption was measured at 4 hours (**A-C**) and 24 hours (**D-F**) timepoints. Statistical analyses are depicted in **Supplementary Table 2**. *p<0.05, **p<0.01. ns: non-significant.

**Supplementary Figure 13.**
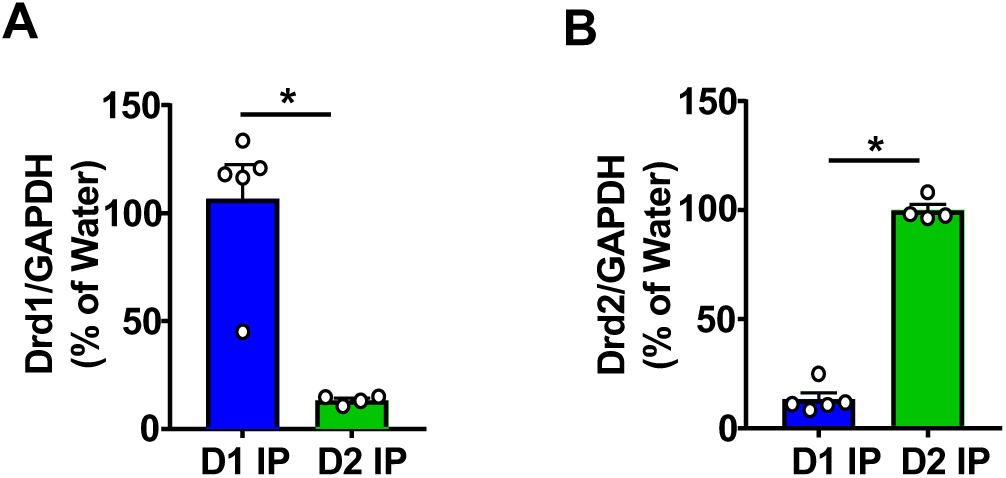
Validation of the RiboTag technique. Polysomal mRNA levels in D1+ vs. D2+ neurons of Drd1 (**A**) and Drd2 (**B**) were determined by RT-qPCR. Data are presented as the average ratio of a transcript to GAPDH±SEM and expressed as % of water control. Statistical analyses are depicted in **Supplementary Table 2**. *p<0.05.

**Supplementary Table 1.** Alcohol drinking averages of mouse groups used in the study. Shown are average alcohol intake +/- SEM of the last 3 24-hours sessions for each of the experimental groups.

| **Alcohol** | **Average Consumption**<br>**(g/kg/24hr)** | **SEM (±g/kg/24hr)** |
| --- | --- | --- |
| **1B-D and 2A-C** | 12.14 | 1.14 |
| **1E-H, 2D-I, S1C-D, S2C-F**<br>**and S4D-I,** | 12.52 | 1.63 |
| **3G-L, S1A, S14A** | 17.89 | 1.47 |
| **4B and S7** | 13.25 | 0.97 |
| **6B** | 14.62 | 0.95 |
| **6C** | 11.30 | 0.87 |
| **7B-G and S12** | 10.88 | 2.62 |
| **S2A-B and S4A-C** | 13.60 | 0.96 |
| **4C-D** | 12.55 | 1.37 |
| **S1B, S5, S14B** | 16.41 | 1.17 |
| **Sucrose** | **Average Consumption**<br>**(ml/kg/24hr)** | **SEM**<br>**(±ml/kg/24hr)** |
| **S13** | 213.56 | 21.07 |

**Supplementary Table 2.**
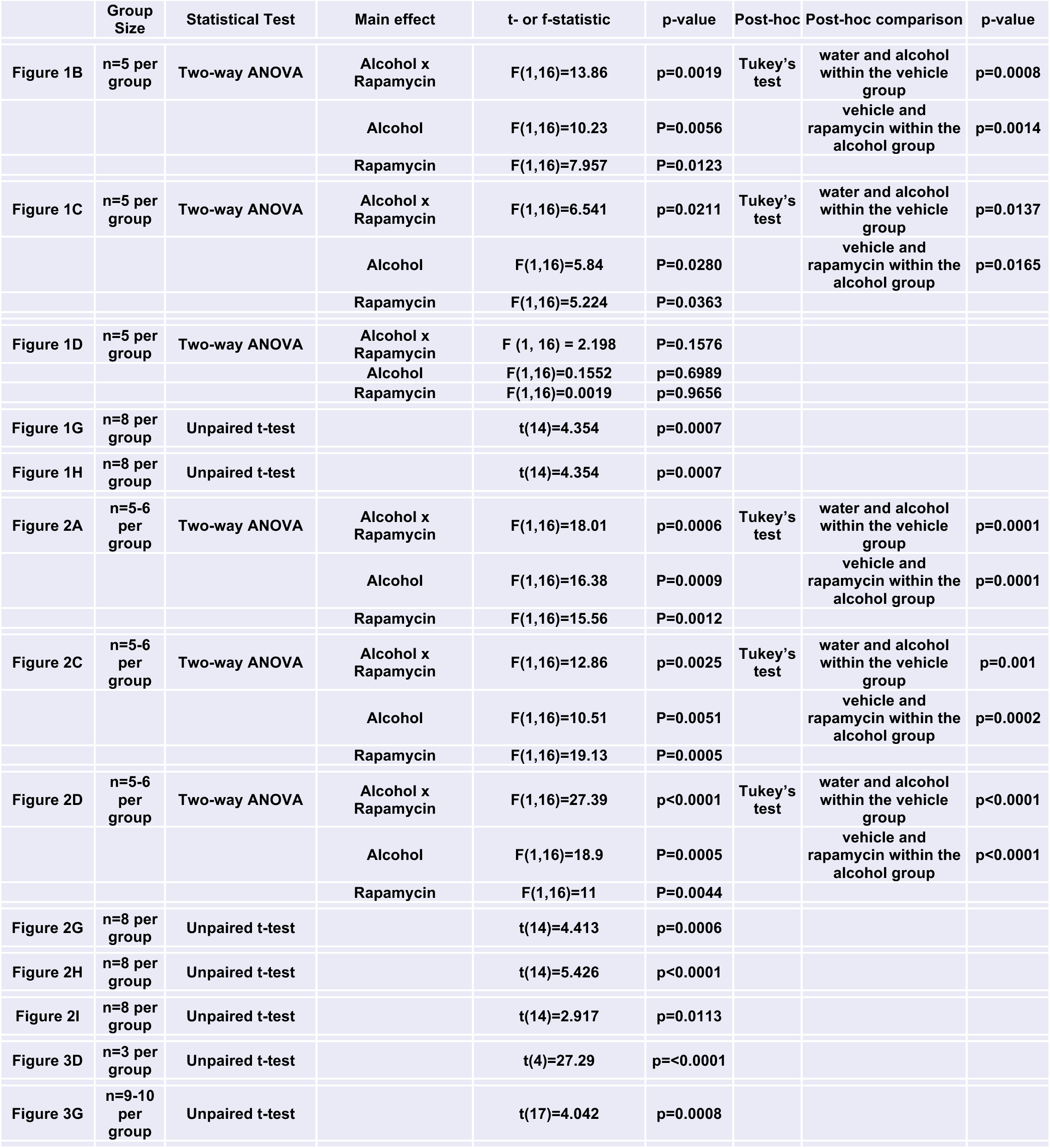
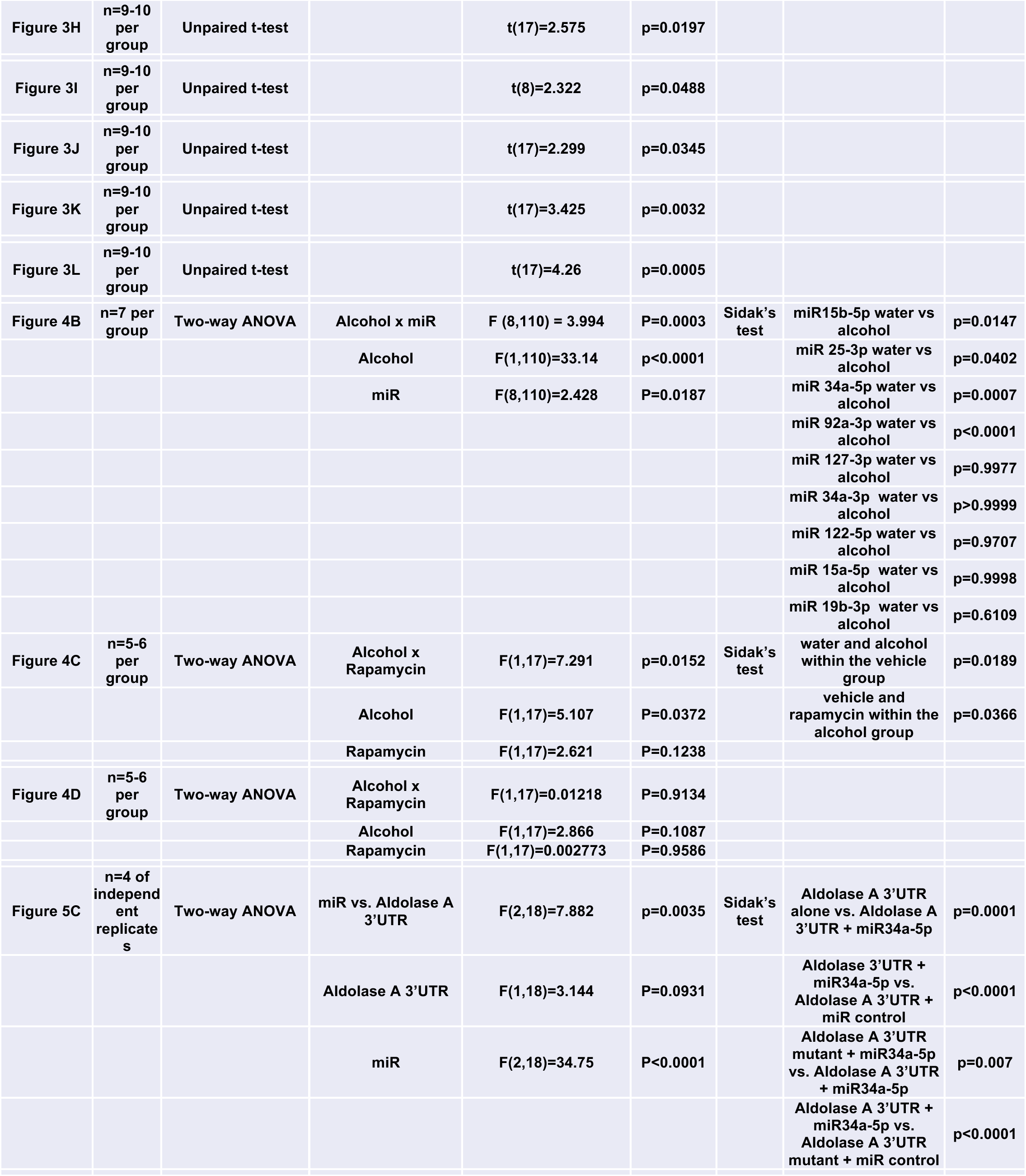
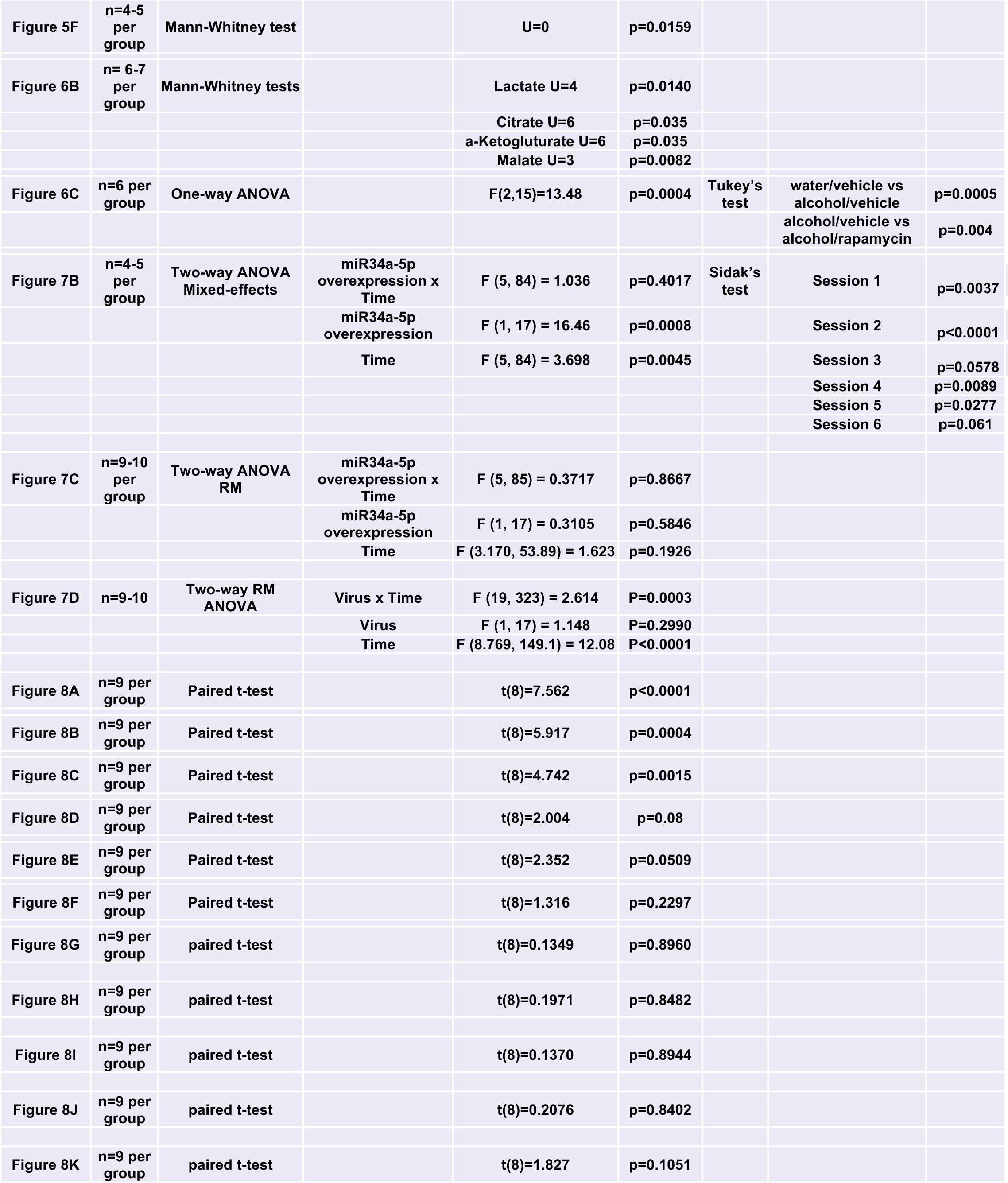
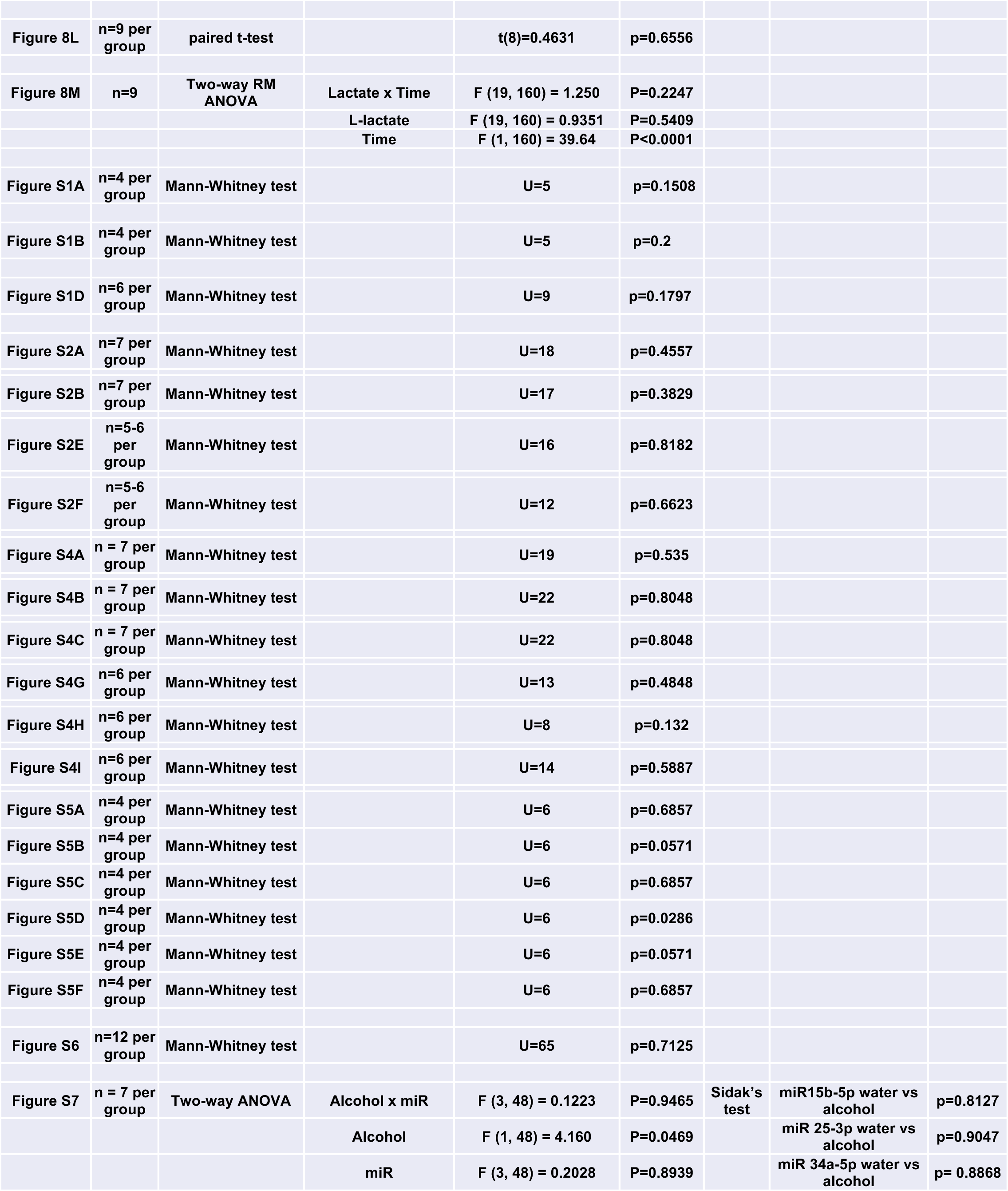
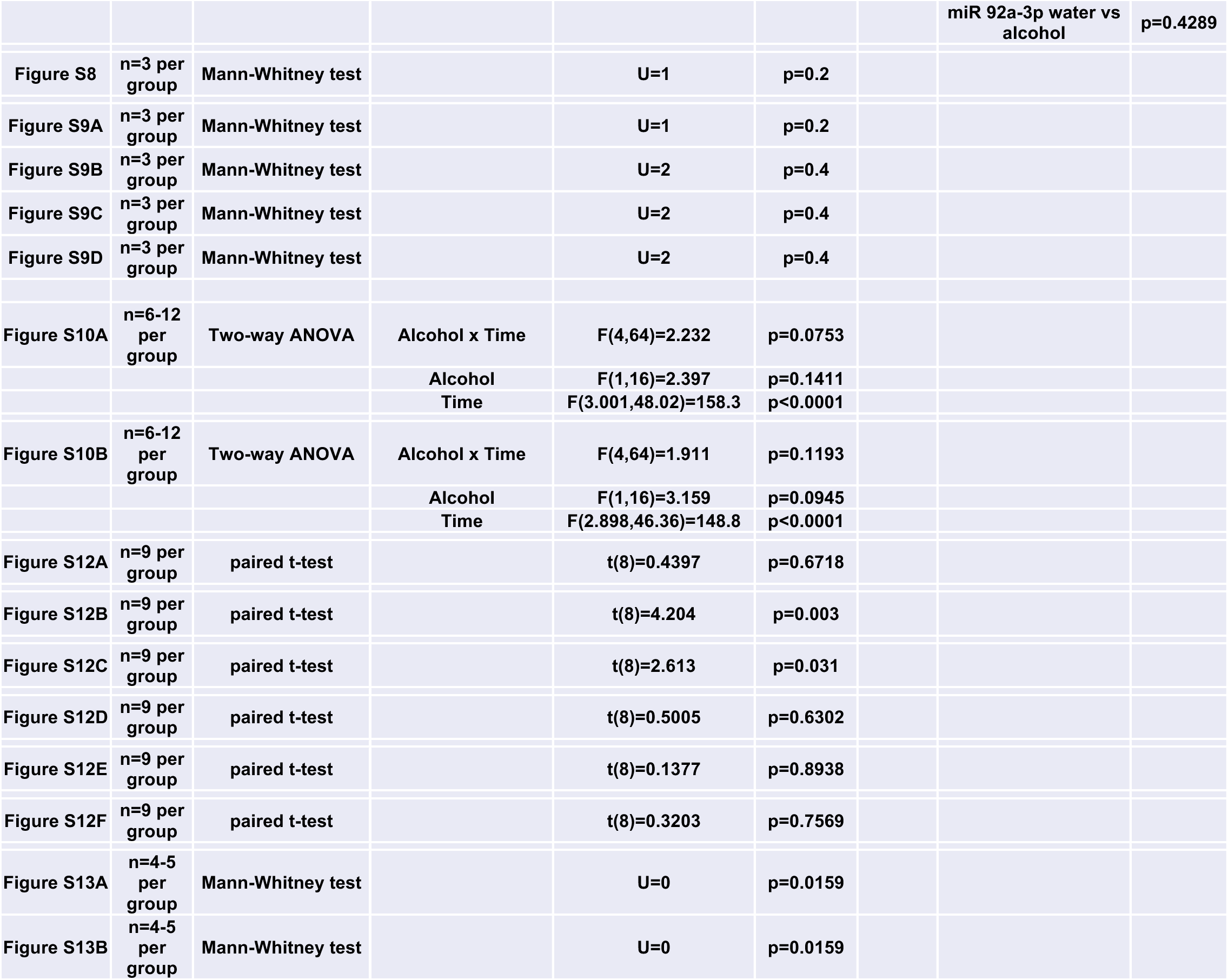
Statistical analyses of the data. Shown are figure number, number of animals or replicates, statistical test and statistical analysis.

**Supplementary Table 3.**
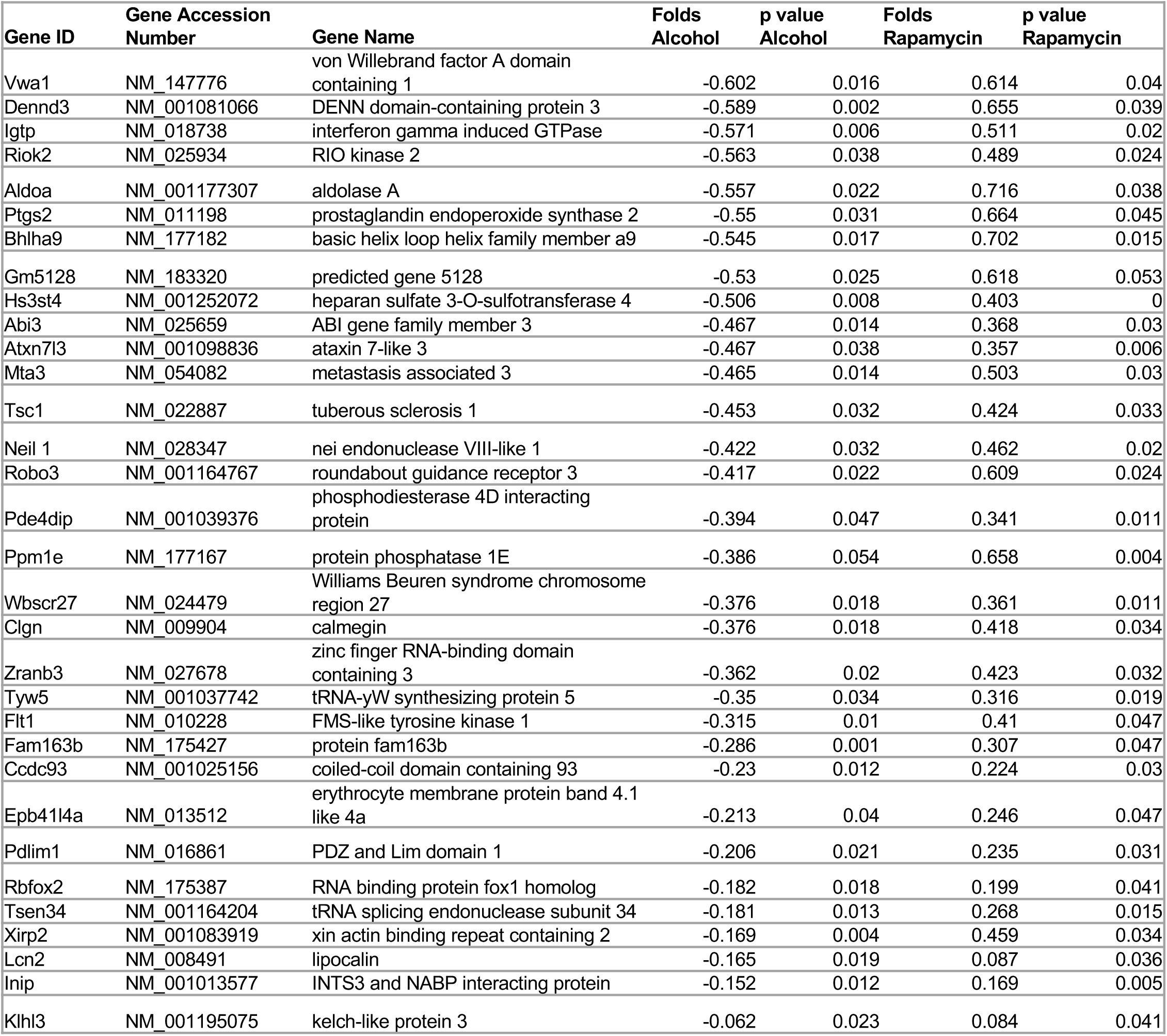
Alcohol decreases the translation of transcripts in an mTORC1 dependent manner. After 7 weeks of IA20%2BC, the NAc of mice was dissected at the end of the 24 hours alcohol withdrawal session. Control animals had access to water only. RNAseq data ^19^ are sorted in ascending order of fold change of alcohol+vehicle divided by water+vehicle with negative values indicating decreased translation by alcohol. Positive fold change of alcohol+vehicle divided by alcohol+rapamycin indicates a reversal of translation attenuation by rapamycin. Student-t-test n = 3.

**Supplementary Table 4.**
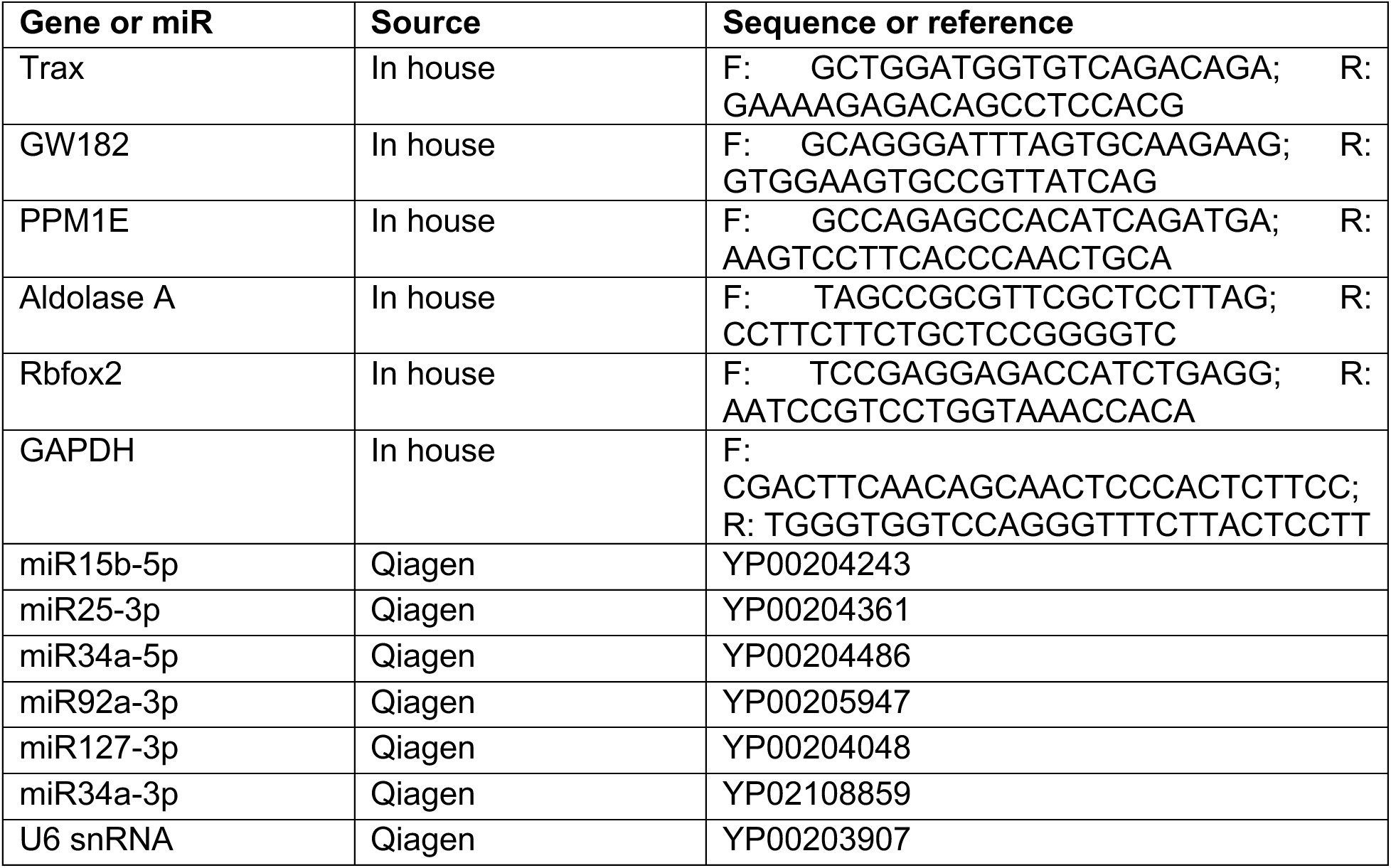
List of primers used in the study. Shown are genes and miR names and corresponding primer information used for RT-qPCR.

**Supplementary Table 5.** List of antibodies used in the study. Shown are antibodies, references, and manufacturers.

| **Antibodies** |  |  |
| --- | --- | --- |
| Rabbit anti-Aldolase A | Cell Signaling | 3188s |
| Rabbit anti-PPM1E | Abnova | PAB21197 |
| Rabbit anti-Rbfox2 | Bethyl | A300-864A |
| Rabbit anti-Trax | Abgent | AP13947a |
| Rabbit anti-GW182 | Sigma | SAB2102506-100UL |
| Mouse IgM anti-Tubulin | Santa Cruz Biotechnology | SC-8035 |
| Mouse anti-GAPDH | Sigma | G8795 |
| Rabbit anti-phospho S6 ribosomal protein | Cell Signaling | 2211s |
| Chicken anti-GFP | Life Technologies | A10262 |
| Guinea pig anti-NeuN | Millipore | ABN90 |
| Donkey anti-rabbit horseradish peroxidase | Jackson ImmunoResearch | 711-035-152 |
| Donkey anti-mouse horseradish peroxidase | Jackson ImmunoResearch | 715-035-150 |
| Goat anti-mouse IgM horseradish peroxidase | Jackson ImmunoResearch | 115-035-020 |
| Donkey anti Rabbit Alexa fluor 488 | Thermo Fisher Scientific | A21206 |
| Goat anti Guinea Pig Alexa fluor 594 | Thermo Fisher Scientific | A11076 |
| Goat anti Chicken Alexa fluor 488 | Thermo Fisher Scientific | A11039 |
| Anti-GFP | Memorial Sloan-Kettering Monoclonal Antibody Facility | clone 19C8 and clone 19F7 |

## References

1 Liu, G. Y. & Sabatini, D. M. mTOR at the nexus of nutrition, growth, ageing and disease. Nat Rev Mol Cell Biol 21, 183–203, doi:10.1038/s41580-019-0199-y (2020).

2 Napolitano, G., Di Malta, C. & Ballabio, A. Non-canonical mTORC1 signaling at the lysosome. Trends Cell Biol 32, 920–931, doi:10.1016/j.tcb.2022.04.012 (2022).

3 Kim, J. & Guan, K. L. mTOR as a central hub of nutrient signalling and cell growth. Nat Cell Biol 21, 63–71, doi:10.1038/s41556-018-0205-1 (2019).

4 Santini, E., Huynh, T. N. & Klann, E. Mechanisms of translation control underlying long-lasting synaptic plasticity and the consolidation of long-term memory. Prog Mol Biol Transl Sci 122, 131–167, doi:10.1016/B978-0-12-420170-5.00005-2 (2014).

5 Lipton, J. O. & Sahin, M. The neurology of mTOR. Neuron 84, 275–291, doi:10.1016/j.neuron.2014.09.034 (2014).

6 Costa-Mattioli, M., Sossin, W. S., Klann, E. & Sonenberg, N. Translational control of long-lasting synaptic plasticity and memory. Neuron 61, 10–26, doi:10.1016/j.neuron.2008.10.055 (2009).

7 Stoica, L. et al. Selective pharmacogenetic inhibition of mammalian target of Rapamycin complex I (mTORC1) blocks long-term synaptic plasticity and memory storage. Proc Natl Acad Sci U S A 108, 3791–3796, doi:10.1073/pnas.1014715108 (2011).

8 Buffington, S. A., Huang, W. & Costa-Mattioli, M. Translational control in synaptic plasticity and cognitive dysfunction. Annu Rev Neurosci 37, 17–38, doi:10.1146/annurev-neuro-071013-014100 (2014).

9 Costa-Mattioli, M. & Monteggia, L. M. mTOR complexes in neurodevelopmental and neuropsychiatric disorders. Nat Neurosci 16, 1537–1543, doi:10.1038/nn.3546 (2013).

10 Neasta, J., Barak, S., Hamida, S. B. & Ron, D. mTOR complex 1: a key player in neuroadaptations induced by drugs of abuse. J Neurochem 130, 172–184, doi:10.1111/jnc.12725 (2014).

11 Ben Hamida, S., et al. The small G protein H-Ras in the mesolimbic system is a molecular gateway to alcohol-seeking and excessive drinking behaviors. J Neurosci 32, 15849–15858, doi:10.1523/JNEUROSCI.2846-12.2012 (2012).

12 Neasta, J., Ben Hamida, S., Yowell, Q. V., Carnicella, S. & Ron, D. AKT signaling pathway in the nucleus accumbens mediates excessive alcohol drinking behaviors. Biol Psychiatry 70, 575–582, doi:10.1016/j.biopsych.2011.03.019 (2011).

13 Manning, B. D. & Cantley, L. C. AKT/PKB signaling: navigating downstream. Cell 129, 1261–1274, doi:10.1016/j.cell.2007.06.009 (2007).

14 Hoeffer, C. A. & Klann, E. mTOR signaling: at the crossroads of plasticity, memory and disease. Trends Neurosci 33, 67–75, doi:10.1016/j.tins.2009.11.003 (2010).

15 Neasta, J., Ben Hamida, S., Yowell, Q., Carnicella, S. & Ron, D. Role for mammalian target of rapamycin complex 1 signaling in neuroadaptations underlying alcohol-related disorders. Proc Natl Acad Sci U S A 107, 20093–20098, doi:10.1073/pnas.1005554107 (2010).

16 Laguesse, S., Morisot, N., Phamluong, K. & Ron, D. Region specific activation of the AKT and mTORC1 pathway in response to excessive alcohol intake in rodents. Addict Biol 22, 1856–1869, doi:10.1111/adb.12464 (2017).

17 Li, J., Kim, S. G. & Blenis, J. Rapamycin: one drug, many effects. Cell Metab 19, 373–379, doi:10.1016/j.cmet.2014.01.001 (2014).

18 Liu, F. et al. mTORC1-dependent translation of collapsin response mediator protein-2 drives neuroadaptations underlying excessive alcohol-drinking behaviors. Mol Psychiatry 22, 89–101, doi:10.1038/mp.2016.12 (2017).

19 Laguesse, S. et al. Prosapip1-Dependent Synaptic Adaptations in the Nucleus Accumbens Drive Alcohol Intake, Seeking, and Reward. Neuron 96, 145–159 e148, doi:10.1016/j.neuron.2017.08.037 (2017).

20 Beckley, J. T. et al. The First Alcohol Drink Triggers mTORC1-Dependent Synaptic Plasticity in Nucleus Accumbens Dopamine D1 Receptor Neurons. J Neurosci 36, 701–713, doi:10.1523/JNEUROSCI.2254-15.2016 (2016).

21 Ben Hamida, S., et al. Mammalian target of rapamycin complex 1 and its downstream effector collapsin response mediator protein-2 drive reinstatement of alcohol reward seeking. Addict Biol 24, 908–920, doi:10.1111/adb.12653 (2019).

22 Ehinger, Y. et al. Brain-specific inhibition of mTORC1 eliminates side effects resulting from mTORC1 blockade in the periphery and reduces alcohol intake in mice. Nat Commun 12, 4407, doi:10.1038/s41467-021-24567-x (2021).

23 Bartel, D. P. MicroRNAs: genomics, biogenesis, mechanism, and function. Cell 116, 281–297, doi:10.1016/s0092-8674(04)00045-5 (2004).

24 Gu, S. & Kay, M. A. How do miRNAs mediate translational repression? Silence 1, 11, doi:10.1186/1758-907X-1-11 (2010).

25 Gebert, L. F. R. & MacRae, I. J. Regulation of microRNA function in animals. Nat Rev Mol Cell Biol 20, 21–37, doi:10.1038/s41580-018-0045-7 (2019).

26 Ha, M. & Kim, V. N. Regulation of microRNA biogenesis. Nat Rev Mol Cell Biol 15, 509–524, doi:10.1038/nrm3838 (2014).

27 Carnicella, S., Ron, D. & Barak, S. Intermittent ethanol access schedule in rats as a preclinical model of alcohol abuse. Alcohol 48, 243–252, doi:10.1016/j.alcohol.2014.01.006 (2014).

28 Penhoet, E. E., Kochman, M. & Rutter, W. J. Ioslation of fructose diphosphate aldolases A, B, and C. Biochemistry 8, 4391–4395, doi:10.1021/bi00839a025 (1969).

29 Voss, M. et al. Ppm1E is an in cellulo AMP-activated protein kinase phosphatase. Cell Signal 23, 114–124, doi:10.1016/j.cellsig.2010.08.010 (2011).

30 Arya, A. D., Wilson, D. I., Baralle, D. & Raponi, M. RBFOX2 protein domains and cellular activities. Biochem Soc Trans 42, 1180–1183, doi:10.1042/BST20140050 (2014).

31 Gerfen, C. R. & Surmeier, D. J. Modulation of striatal projection systems by dopamine. Annu Rev Neurosci 34, 441–466, doi:10.1146/annurev-neuro-061010-113641 (2011).

32 Lesiak, A. J. & Neumaier, J. F. RiboTag: Not Lost in Translation. Neuropsychopharmacology 41, 374–376, doi:10.1038/npp.2015.262 (2016).

33 Wachsmuth, E. D., Thoner, M. & Pfleiderer, G. The cellular distribution of aldolase isozymes in rat kidney and brain determined in tissue sections by the immuno-histochemical method. Histochemistry 45, 143–161, doi:10.1007/BF00495158 (1975).

34 Berardini, T. Z., Drygas-Williams, M., Callard, G. V. & Tolan, D. R. Identification of neuronal isozyme specific residues by comparison of goldfish aldolase C to other aldolases. Comp Biochem Physiol A Physiol 117, 471–476, doi:10.1016/s0300-9629(96)00396-9 (1997).

35 Thompson, R. J., Kynoch, P. A. & Willson, V. J. Cellular localization of aldolase C subunits in human brain. Brain Res 232, 489–493, doi:10.1016/0006-8993(82)90294-3 (1982).

36 Magistretti, P. J. & Allaman, I. Lactate in the brain: from metabolic end-product to signalling molecule. Nat Rev Neurosci 19, 235–249, doi:10.1038/nrn.2018.19 (2018).

37 Dienel, G. A. Brain Glucose Metabolism: Integration of Energetics with Function. Physiol Rev 99, 949–1045, doi:10.1152/physrev.00062.2017 (2019).

38 Ter Horst, K. W., et al. Striatal dopamine regulates systemic glucose metabolism in humans and mice. Sci Transl Med 10, doi:10.1126/scitranslmed.aar3752 (2018).

39 Alhadeff, A. L. et al. Natural and Drug Rewards Engage Distinct Pathways that Converge on Coordinated Hypothalamic and Reward Circuits. Neuron 103, 891–908 e896, doi:10.1016/j.neuron.2019.05.050 (2019).

40 Nall, R. W., Heinsbroek, J. A., Nentwig, T. B., Kalivas, P. W. & Bobadilla, A. C. Circuit selectivity in drug versus natural reward seeking behaviors. J Neurochem 157, 1450–1472, doi:10.1111/jnc.15297 (2021).

41 Lund, J. et al. The anorectic and thermogenic effects of pharmacological lactate in male mice are confounded by treatment osmolarity and co-administered counterions. Nat Metab 5, 677–698, doi:10.1038/s42255-023-00780-4 (2023).

42 Cozzoli, D. K. et al. Functional regulation of PI3K-associated signaling in the accumbens by binge alcohol drinking in male but not female mice. Neuropharmacology 105, 164–174, doi:10.1016/j.neuropharm.2016.01.010 (2016).

43 Ehinger, Y., Phamluong, K. & Ron, D. Sex differences in the interaction between alcohol and mTORC1. bioRxiv, 2023.2010.2004.560781, doi:10.1101/2023.10.04.560781 (2023).

44 Valentino, R. J., Van Bockstaele, E. & Bangasser, D. Sex-specific cell signaling: the corticotropin-releasing factor receptor model. Trends Pharmacol Sci 34, 437–444, doi:10.1016/j.tips.2013.06.004 (2013).

45 Marrocco, J. & McEwen, B. S. Sex in the brain: hormones and sex differences. Dialogues Clin Neurosci 18, 373–383, doi:10.31887/DCNS.2016.18.4/jmarrocco (2016).

46 Sneddon, E. A. et al. Sex chromosome and gonadal hormone contributions to binge-like and aversion-resistant ethanol drinking behaviors in Four Core Genotypes mice. Front Psychiatry 14, 1098387, doi:10.3389/fpsyt.2023.1098387 (2023).

47 Sneddon, E. A. et al. Gonadal hormones and sex chromosome complement differentially contribute to ethanol intake, preference, and relapse-like behaviour in four core genotypes mice. Addict Biol 27, e13222, doi:10.1111/adb.13222 (2022).

48 Hwa, L., Besheer, J. & Kash, T. Glutamate plasticity woven through the progression to alcohol use disorder: a multi-circuit perspective. F1000Res 6, 298, doi:10.12688/f1000research.9609.1 (2017).

49 Pati, D., Kelly, K., Stennett, B., Frazier, C. J. & Knackstedt, L. A. Alcohol consumption increases basal extracellular glutamate in the nucleus accumbens core of Sprague-Dawley rats without increasing spontaneous glutamate release. Eur J Neurosci 44, 1896–1905, doi:10.1111/ejn.13284 (2016).

50 Bockaert, J. & Marin, P. mTOR in Brain Physiology and Pathologies. Physiol Rev 95, 1157–1187, doi:10.1152/physrev.00038.2014 (2015).

51 Morisot, N. et al. mTORC1 in the orbitofrontal cortex promotes habitual alcohol seeking. Elife 8, e51333, doi:10.7554/eLife.51333 (2019).

52 Wang, J. et al. Ethanol induces long-term facilitation of NR2B-NMDA receptor activity in the dorsal striatum: implications for alcohol drinking behavior. J Neurosci 27, 3593–3602, doi:10.1523/JNEUROSCI.4749-06.2007 (2007).

53 Lovinger, D. M., White, G. & Weight, F. F. Ethanol inhibits NMDA-activated ion current in hippocampal neurons. Science 243, 1721–1724, doi:10.1126/science.2467382 (1989).

54 Li, N. et al. mTOR-dependent synapse formation underlies the rapid antidepressant effects of NMDA antagonists. Science 329, 959–964, doi:10.1126/science.1190287 (2010).

55 Workman, E. R., Niere, F. & Raab-Graham, K. F. mTORC1-dependent protein synthesis underlying rapid antidepressant effect requires GABABR signaling. Neuropharmacology 73, 192–203, doi:10.1016/j.neuropharm.2013.05.037 (2013).

56 Krystal, J. H., Kavalali, E. T. & Monteggia, L. M. Ketamine and rapid antidepressant action: new treatments and novel synaptic signaling mechanisms. Neuropsychopharmacology 49, 41–50, doi:10.1038/s41386-023-01629-w (2024).

57 Asada, K. et al. Rescuing dicer defects via inhibition of an anti-dicing nuclease. Cell Rep 9, 1471–1481, doi:10.1016/j.celrep.2014.10.021 (2014).

58 Baraban, J. M., Shah, A. & Fu, X. Multiple Pathways Mediate MicroRNA Degradation: Focus on the Translin/Trax RNase Complex. Adv Pharmacol 82, 1–20, doi:10.1016/bs.apha.2017.08.003 (2018).

59 Chen, Y. et al. A DDX6-CNOT1 complex and W-binding pockets in CNOT9 reveal direct links between miRNA target recognition and silencing. Mol Cell 54, 737–750, doi:10.1016/j.molcel.2014.03.034 (2014).

60 Ogorek, B. et al. TSC2 regulates microRNA biogenesis via mTORC1 and GSK3beta. Hum Mol Genet 27, 1654–1663, doi:10.1093/hmg/ddy073 (2018).

61 Niere, F. et al. Aberrant DJ-1 expression underlies L-type calcium channel hypoactivity in dendrites in tuberous sclerosis complex and Alzheimer’s disease. Proc Natl Acad Sci U S A 120, e2301534120, doi:10.1073/pnas.2301534120 (2023).

62 Niere, F. et al. Analysis of Proteins That Rapidly Change Upon Mechanistic/Mammalian Target of Rapamycin Complex 1 (mTORC1) Repression Identifies Parkinson Protein 7 (PARK7) as a Novel Protein Aberrantly Expressed in Tuberous Sclerosis Complex (TSC). Mol Cell Proteomics 15, 426–444, doi:10.1074/mcp.M115.055079 (2016).

63 Lam, H. C. et al. Rapamycin-induced miR-21 promotes mitochondrial homeostasis and adaptation in mTORC1 activated cells. Oncotarget 8, 64714–64727, doi:10.18632/oncotarget.19947 (2017).

64 Kye, M. J. et al. SMN regulates axonal local translation via miR-183/mTOR pathway. Hum Mol Genet 23, 6318–6331, doi:10.1093/hmg/ddu350 (2014).

65 Kar, A. N. et al. MicroRNAs 21 and 199a-3p Regulate Axon Growth Potential through Modulation of Pten and mTor mRNAs. eNeuro 8, doi:10.1523/ENEURO.0155-21.2021 (2021).

66 Zhang, Y., Huang, B., Wang, H. Y., Chang, A. & Zheng, X. F. S. Emerging Role of MicroRNAs in mTOR Signaling. Cell Mol Life Sci 74, 2613–2625, doi:10.1007/s00018-017-2485-1 (2017).

67 Bazrgar, M., Khodabakhsh, P., Prudencio, M., Mohagheghi, F. & Ahmadiani, A. The role of microRNA-34 family in Alzheimer’s disease: A potential molecular link between neurodegeneration and metabolic disorders. Pharmacol Res 172, 105805, doi:10.1016/j.phrs.2021.105805 (2021).

68 Haramati, S. et al. MicroRNA as repressors of stress-induced anxiety: the case of amygdalar miR-34. J Neurosci 31, 14191–14203, doi:10.1523/JNEUROSCI.1673-11.2011 (2011).

69 Murphy, C. P. & Singewald, N. Role of MicroRNAs in Anxiety and Anxiety-Related Disorders. Curr Top Behav Neurosci 42, 185–219, doi:10.1007/7854_2019_109 (2019).

70 Lai, Y. W. et al. Drosophila microRNA-34 Impairs Axon Pruning of Mushroom Body gamma Neurons by Downregulating the Expression of Ecdysone Receptor. Sci Rep 6, 39141, doi:10.1038/srep39141 (2016).

71 Malmevik, J. et al. Distinct cognitive effects and underlying transcriptome changes upon inhibition of individual miRNAs in hippocampal neurons. Sci Rep 6, 19879, doi:10.1038/srep19879 (2016).

72 McNeill, E. M. et al. The conserved microRNA miR-34 regulates synaptogenesis via coordination of distinct mechanisms in presynaptic and postsynaptic cells. Nat Commun 11, 1092, doi:10.1038/s41467-020-14761-8 (2020).

73 Dias, B. G. et al. Amygdala-dependent fear memory consolidation via miR-34a and Notch signaling. Neuron 83, 906–918, doi:10.1016/j.neuron.2014.07.019 (2014).

74 Raab-Graham, K. F., Haddick, P. C., Jan, Y. N. & Jan, L. Y. Activity- and mTOR-dependent suppression of Kv1.1 channel mRNA translation in dendrites. Science 314, 144–148, doi:10.1126/science.1131693 (2006).

75 Sosanya, N. M. et al. Degradation of high affinity HuD targets releases Kv1.1 mRNA from miR-129 repression by mTORC1. J Cell Biol 202, 53–69, doi:10.1083/jcb.201212089 (2013).

76 Sosanya, N. M., Brager, D. H., Wolfe, S., Niere, F. & Raab-Graham, K. F. Rapamycin reveals an mTOR-independent repression of Kv1.1 expression during epileptogenesis. Neurobiol Dis 73, 96–105, doi:10.1016/j.nbd.2014.09.011 (2015).

77 Pirovich, D. B., Da’dara, A. A. & Skelly, P. J. Multifunctional Fructose 1,6-Bisphosphate Aldolase as a Therapeutic Target. Front Mol Biosci 8, 719678, doi:10.3389/fmolb.2021.719678 (2021).

78 Volkow, N. D. et al. Decreased brain metabolism in neurologically intact healthy alcoholics. Am J Psychiatry 149, 1016–1022, doi:10.1176/ajp.149.8.1016 (1992).

79 Volkow, N. D. et al. Low doses of alcohol substantially decrease glucose metabolism in the human brain. Neuroimage 29, 295–301, doi:10.1016/j.neuroimage.2005.07.004 (2006).

80 Volkow, N. D. et al. Alcohol decreases baseline brain glucose metabolism more in heavy drinkers than controls but has no effect on stimulation-induced metabolic increases. J Neurosci 35, 3248–3255, doi:10.1523/JNEUROSCI.4877-14.2015 (2015).

81 Tomasi, D., Wang, G. J. & Volkow, N. D. Energetic cost of brain functional connectivity. Proc Natl Acad Sci U S A 110, 13642–13647, doi:10.1073/pnas.1303346110 (2013).

82 Wang, G. J. et al. Alcohol intoxication induces greater reductions in brain metabolism in male than in female subjects. Alcohol Clin Exp Res 27, 909–917, doi:10.1097/01.ALC.0000071740.56375.BA (2003).

83 Dutchak, P. A. et al. Loss of a Negative Regulator of mTORC1 Induces Aerobic Glycolysis and Altered Fiber Composition in Skeletal Muscle. Cell Rep 23, 1907–1914, doi:10.1016/j.celrep.2018.04.058 (2018).

84 Kim, B. et al. Cytoplasmic zinc promotes IL-1beta production by monocytes and macrophages through mTORC1-induced glycolysis in rheumatoid arthritis. Sci Signal 15, eabi7400, doi:10.1126/scisignal.abi7400 (2022).

85 Sun, Q. et al. Mammalian target of rapamycin up-regulation of pyruvate kinase isoenzyme type M2 is critical for aerobic glycolysis and tumor growth. Proc Natl Acad Sci U S A 108, 4129–4134, doi:10.1073/pnas.1014769108 (2011).

86 Xu, Y. et al. Asparagine reinforces mTORC1 signaling to boost thermogenesis and glycolysis in adipose tissues. EMBO J 40, e108069, doi:10.15252/embj.2021108069 (2021).

87 Xu, L., Nan, J. & Lan, Y. The Nucleus Accumbens: A Common Target in the Comorbidity of Depression and Addiction. Front Neural Circuits 14, 37, doi:10.3389/fncir.2020.00037 (2020).

88 Faria-Pereira, A. & Morais, V. A. Synapses: The Brain’s Energy-Demanding Sites. Int J Mol Sci 23, doi:10.3390/ijms23073627 (2022).

89 Saxton, R. A. & Sabatini, D. M. mTOR Signaling in Growth, Metabolism, and Disease. Cell 168, 960–976, doi:10.1016/j.cell.2017.02.004 (2017).

90 Panov, A., Orynbayeva, Z., Vavilin, V. & Lyakhovich, V. Fatty acids in energy metabolism of the central nervous system. Biomed Res Int 2014, 472459, doi:10.1155/2014/472459 (2014).

91 Wlaz, P. et al. Acute anticonvulsant effects of capric acid in seizure tests in mice. Prog Neuropsychopharmacol Biol Psychiatry 57, 110–116, doi:10.1016/j.pnpbp.2014.10.013 (2015).

92 Ebert, D., Haller, R. G. & Walton, M. E. Energy Contribution of Octanoate to Intact Rat Brain Metabolism Measured by ^13^C Nuclear Magnetic Resonance Spectroscopy. The Journal of Neuroscience 23, 5928–5935, doi:10.1523/jneurosci.23-13-05928.2003 (2003).

93 Jiang, L. et al. Increased brain uptake and oxidation of acetate in heavy drinkers. J Clin Invest 123, 1605–1614, doi:10.1172/JCI65153 (2013).

94 Volkow, N. D. et al. Acute alcohol intoxication decreases glucose metabolism but increases acetate uptake in the human brain. Neuroimage 64, 277–283, doi:10.1016/j.neuroimage.2012.08.057 (2013).

95 Jin, S. et al. Brain ethanol metabolism by astrocytic ALDH2 drives the behavioural effects of ethanol intoxication. Nat Metab 3, 337–351, doi:10.1038/s42255-021-00357-z (2021).

96 Dienel, G. A. & Hertz, L. Glucose and lactate metabolism during brain activation. J Neurosci Res 66, 824–838, doi:10.1002/jnr.10079 (2001).

97 Machler, P. et al. In Vivo Evidence for a Lactate Gradient from Astrocytes to Neurons. Cell Metab 23, 94–102, doi:10.1016/j.cmet.2015.10.010 (2016).

98 Dienel, G. A. Lack of appropriate Stoichiometry: Strong evidence against an energetically importand astrocytes-neuron lactate shuttle in the brain. Journal of Neuroscience Research, 2103 (2017).

99 Gjedde, A. & Marrett, S. Glycolysis in neurons, not astrocytes, delays oxidative metabolism of human visual cortex during sustained checkerboard stimulation in vivo. J Cereb Blood Flow Metab 21, 1384–1392, doi:10.1097/00004647-200112000-00002 (2001).

100 Cisternas, P., Salazar, P., Silva-Alvarez, C., Barros, L. F. & Inestrosa, N. C. Activation of Wnt Signaling in Cortical Neurons Enhances Glucose Utilization through Glycolysis. J Biol Chem 291, 25950–25964, doi:10.1074/jbc.M116.735373 (2016).

101 Hollnagel, J. O. et al. Lactate Attenuates Synaptic Transmission and Affects Brain Rhythms Featuring High Energy Expenditure. iScience 23, 101316, doi:10.1016/j.isci.2020.101316 (2020).

102 Li, H. et al. Neurons require glucose uptake and glycolysis in vivo. Cell Rep 42, 112335, doi:10.1016/j.celrep.2023.112335 (2023).

103 Diaz-Garcia, C. M. et al. Neuronal Stimulation Triggers Neuronal Glycolysis and Not Lactate Uptake. Cell Metab 26, 361–374 e364, doi:10.1016/j.cmet.2017.06.021 (2017).

104 Dai, X., Lv, X., Thompson, E. W. & Ostrikov, K. K. Histone lactylation: epigenetic mark of glycolytic switch. Trends Genet 38, 124–127, doi:10.1016/j.tig.2021.09.009 (2022).

105 de Castro Abrantes, H., et al. The Lactate Receptor HCAR1 Modulates Neuronal Network Activity through the Activation of G(alpha) and G(betagamma) Subunits. J Neurosci 39, 4422–4433, doi:10.1523/JNEUROSCI.2092-18.2019 (2019).

106 Zhang, D. et al. Metabolic regulation of gene expression by histone lactylation. Nature 574, 575–580, doi:10.1038/s41586-019-1678-1 (2019).

107 Hagihara, H. et al. Protein lactylation induced by neural excitation. Cell Rep 37, 109820, doi:10.1016/j.celrep.2021.109820 (2021).

108 Li, X. et al. Lactate metabolism in human health and disease. Signal Transduct Target Ther 7, 305, doi:10.1038/s41392-022-01151-3 (2022).

109 Dencker, D. et al. Ketogenic Diet Suppresses Alcohol Withdrawal Syndrome in Rats. Alcohol Clin Exp Res 42, 270–277, doi:10.1111/acer.13560 (2018).

110 Wiers, C. E. et al. Ketogenic diet reduces alcohol withdrawal symptoms in humans and alcohol intake in rodents. Sci Adv 7, doi:10.1126/sciadv.abf6780 (2021).

111 Bornebusch, A. B. et al. Effects of ketogenic diet and ketone monoester supplement on acute alcohol withdrawal symptoms in male mice. Psychopharmacology (Berl) 238, 833–844, doi:10.1007/s00213-020-05735-1 (2021).

112 Willemse, L., Terburgh, K. & Louw, R. A ketogenic diet alters mTOR activity, systemic metabolism and potentially prevents collagen degradation associated with chronic alcohol consumption in mice. Metabolomics 19, 43, doi:10.1007/s11306-023-02006-w (2023).

113 McDaniel, S. S., Rensing, N. R., Thio, L. L., Yamada, K. A. & Wong, M. The ketogenic diet inhibits the mammalian target of rapamycin (mTOR) pathway. Epilepsia 52, e7–11, doi:10.1111/j.1528-1167.2011.02981.x (2011).

114 Carrard, A. et al. Peripheral administration of lactate produces antidepressant-like effects. Mol Psychiatry 23, 488, doi:10.1038/mp.2016.237 (2018).

115 Cherix, A. et al. Deletion of Crtc1 leads to hippocampal neuroenergetic impairments associated with depressive-like behavior. Mol Psychiatry, doi:10.1038/s41380-022-01791-5 (2022).

116 Carrard, A. et al. Role of adult hippocampal neurogenesis in the antidepressant actions of lactate. Mol Psychiatry 26, 6723–6735, doi:10.1038/s41380-021-01122-0 (2021).

117 Zhou, P. et al. Interrogating translational efficiency and lineage-specific transcriptomes using ribosome affinity purification. Proc Natl Acad Sci U S A 110, 15395–15400, doi:10.1073/pnas.1304124110 (2013).

118 Agarwal, V., Bell, G. W., Nam, J. W. & Bartel, D. P. Predicting effective microRNA target sites in mammalian mRNAs. Elife 4, e05005, doi:10.7554/eLife.05005 (2015).

119 Sticht, C., De La Torre, C., Parveen, A. & Gretz, N. miRWalk: An online resource for prediction of microRNA binding sites. PLoS One 13, e0206239, doi:10.1371/journal.pone.0206239 (2018).

120 Liu, W. & Wang, X. Prediction of functional microRNA targets by integrative modeling of microRNA binding and target expression data. Genome Biol 20, 18, doi:10.1186/s13059-019-1629-z (2019).

121 Chen, Y. & Wang, X. miRDB: an online database for prediction of functional microRNA targets. Nucleic Acids Res 48, D127–D131, doi:10.1093/nar/gkz757 (2020).

122 Ding, J., Li, X. & Hu, H. TarPmiR: a new approach for microRNA target site prediction. Bioinformatics 32, 2768–2775, doi:10.1093/bioinformatics/btw318 (2016).

123 Darcq, E. et al. MicroRNA-30a-5p in the prefrontal cortex controls the transition from moderate to excessive alcohol consumption. Mol Psychiatry 20, 1261, doi:10.1038/mp.2014.155 (2015).

124 Heiman, M., Kulicke, R., Fenster, R. J., Greengard, P. & Heintz, N. Cell type-specific mRNA purification by translating ribosome affinity purification (TRAP). Nat Protoc 9, 1282–1291, doi:10.1038/nprot.2014.085 (2014).

125 Warnault, V. et al. The BDNF Valine 68 to Methionine Polymorphism Increases Compulsive Alcohol Drinking in Mice That Is Reversed by Tropomyosin Receptor Kinase B Activation. Biol Psychiatry 79, 463–473, doi:10.1016/j.biopsych.2015.06.007 (2016).

